# Transcriptome Analysis Identifies An ASD-Like Phenotype In Oligodendrocytes And Microglia From C58/J Amygdala That Is Dependent On Sex and Sociability

**DOI:** 10.1101/2024.01.15.575733

**Authors:** George D. Dalton, Stephen K. Siecinski, Viktoriya D. Nikolova, Gary P. Cofer, Kathryn Hornburg, Yi Qi, G. Allan Johnson, Yong-Hui Jiang, Sheryl S. Moy, Simon G. Gregory

**Affiliations:** Duke Molecular Physiology Institute, Duke University School of Medicine, Durham, NC USA 27701; Department of Neurology, Duke University School of Medicine,g Durham, NC, USA 27710; Department of Psychiatry, The University of North Carolina at Chapel Hill, Chapel Hill, NC, USA 27516; Center for In Vivo Microscopy, Duke University, Durham, NC 27710; Department of Genetics, Neuroscience, and Pediatrics, Yale University School of Medicine, New Haven, CT, USA 06520

**Keywords:** Autism, oligodendrocyte, microglia, myelin, oxytocin, amygdala, brain, neurodegenerative disease, neuroinflammation, glia

## Abstract

**Background:** Autism Spectrum Disorder (ASD) is a group of neurodevelopmental disorders with higher incidence in males and is characterized by atypical verbal/nonverbal communication, restricted interests that can be accompanied by repetitive behavior, and disturbances in social behavior. This study investigated brain mechanisms that contribute to sociability deficits and sex differences in an ASD animal model.

**Methods:** Sociability was measured in C58/J and C57BL/6J mice using the 3-chamber social choice test. Bulk RNA-Seq and snRNA-Seq identified transcriptional changes in C58/J and C57BL/6J amygdala within which DMRseq was used to measure differentially methylated regions in amygdala.

**Results:** C58/J mice displayed divergent social strata in the 3-chamber test. Transcriptional and pathway signatures revealed immune-related biological processes differ between C58/J and C57BL/6J amygdala. Hypermethylated and hypomethylated genes were identified in C58/J versus C57BL/6J amygdala. snRNA-Seq data in C58/J amygdala identified differential transcriptional signatures within oligodendrocytes and microglia characterized by increased ASD risk gene expression and predicted impaired myelination that was dependent on sex and sociability. RNA velocity, gene regulatory network, and cell communication analysis showed diminished oligodendrocyte/microglia differentiation. Findings were verified using bulk RNA-Seq and demonstrated oxytocin’s beneficial effects on myelin gene expression.

**Limitations:** Our findings are significant. However, limitations can be noted. The cellular mechanisms linking reduced oligodendrocyte differentiation and reduced myelination to an ASD phenotype in C58/J mice need further investigation. Additional snRNA-Seq and spatial studies would determine if effects in oligodendrocytes/microglia are unique to amygdala or if this occurs in other brain regions. Oxytocin’s effects need further examination to understand its potential as an ASD therapeutic.

**Conclusions:** Our work demonstrates the C58/J mouse model’s utility in evaluating the influence of sex and sociability on the transcriptome in concomitant brain regions involved in ASD. Our single-nucleus transcriptome analysis elucidates potential pathological roles of oligodendrocytes and microglia in ASD. This investigation provides details regarding regulatory features disrupted in these cell types, including transcriptional gene dysregulation, aberrant cell differentiation, altered gene regulatory networks, and changes to key pathways that promote microglia/oligodendrocyte differentiation. Our studies provide insight into interactions between genetic risk and epigenetic processes associated with divergent affiliative behavior and lack of positive sociability.

## Background

In 1943, Leo Kanner identified the core symptoms of Autism Spectrum Disorder (ASD) in young children (1). Today, ASD is considered a highly heterogenous and lifelong neurodevelopmental disorder with onset in infancy or early childhood and diagnostic symptoms that include deficits in social communication and interaction, restricted interests and repetitive behaviors, sensory anomalies, and intellectual disability (2). The incidence of ASD in the United States is estimated at 1 in 36 children with a 4:1 male-to-female ratio (3). ASD is a group of disorders characterized by multifactor causation with genetic and non-genetic components. Studies have identified over 1,000 genes that could contribute to ASD risk, as well as chromosomal aberrations, genetic syndromes, metabolic disturbances (mitochondrial dysfunction), epigenetics, and environmental factors (4-6). Despite the identification of possible underlying mechanisms of ASD, a cohesive model of ASD causation does not currently exist and is a pressing unmet need. Furthermore, behavioral interventions remain the standard of care for ASD with no pharmacological treatments available to address ASD core symptoms. Currently antipsychotic drugs (e.g., aripirazole, risperidone) are used to alleviate ASD irritability, but these drugs are not consistently effective and have many adverse side effects (7, 8).

There are several potential targeted treatments for ASD including the use of oxytocin which has been explored as a possible ASD therapeutic (9, 10). Oxytocin is a neuropeptide that is produced in the hypothalamus and secreted both in the blood circulation as a hormone and in the brain where it acts as a neurotransmitter at neuronal oxytocin receptors (11, 12). In rodents, brain oxytocin receptor expression is both developmentally regulated and gender-specific with males having higher expression in regions such as medial amygdala and the hippocampal CA1 region which mediate core behaviors impaired in autism (13, 14). Modulation of the oxytocin pathway may be relevant to ASD because the neuropeptide endogenously mediates a range of social behavior. Disruptions of oxytocin signaling has been shown to result in impaired sociability and communication and leads to repetitive behaviors in ASD animal models and human patients (12, 15). In mouse ASD models, oxytocin treatment improved social deficits, social interaction, and social preference (16, 17). However, clinical studies investigating the efficacy of oxytocin to improve sociability in people with ASD, including our own, have yielded mixed results (10, 18). The failure of oxytocin to act consistently as an ASD therapeutic could be attributed to a lack of understanding about its function. Our current study provides data that helps elucidate the function of oxytocin in a region of the brain that is associated with a core deficit of ASD.

The lack of pharmacological treatments with proven efficacy for ASD is partly because the pathophysiology of ASD remains largely unclear. Abnormal brain growth and structural dissimilarities between ASD and normal brain appear to be important in understanding the symptoms and neuropathology of ASD (19-23). Alterations of brain morphology in ASD patients are considered by some to be due to brain hypoconnectivity/hyperconnectivity, which leads to imbalances in neuronal excitation/inhibition (E/I) and abnormal brain development and function (24-29). In support of these hypotheses, studies suggest that ASD anatomical/functional abnormalities are due to myelination deficits, fewer axons, and oxidative stress (27, 30, 31). However, the specific cellular structures that carry out these processes have still not been adequately identified. Glial cells (oligodendrocytes, astrocytes, microglia) are candidates that may play a role in brain anatomy and functional changes in ASD. They constitute the most abundant cell types in the brain and critically protect the brain’s health under homeostatic conditions. However, in certain pathological states microglia become reactive and secrete proinflammatory cytokines that cause oxidative stress, brain inflammation, neuronal and oligodendrocyte cell death, impaired myelination, neural hypoconnectivity, and dysfunctional synaptic plasticity (32). Like microglia, astrocytes can also become reactive and release reactive oxygen species (ROS) that damage oligodendrocytes and neurons and cause changes in glutamate transport that results in E/I imbalance that is characteristic of ASD (32). Thus, reactive glia can contribute to abnormal cytokine profiles in ASD patients as well as altered brain myelination and white matter density. Importantly, alterations in white matter density in brain regions (amygdala, frontal cortex) that play important roles with ASD core symptoms like sociability have been documented in ASD patients (33, 34). In this study, we attempt to identify the cellular structures that are responsible for ASD pathophysiology as it relates to divergent sociability, and we specifically address the role that glial cells (oligodendrocytes, microglia) and myelination defects play in this process within the amygdala.

Mouse models play an important role in understanding the causes of ASD (35). C58/J is an inbred mouse strain that exhibits low sociability primarily in males and deficits in social transmission of food preference (36, 37). In the 3-chamber social choice test used to assess sociability, our research group has found that C58/J mice exhibit divergent phenotypes with approximately 50% of mice exhibiting positive sociability and 50% exhibiting social avoidance (37). C58/J mice also develop motor stereotypic behaviors including backflipping, “jackhammer” jumping, and upright scrabbling which could reflect abnormal repetitive and social behavior in ASD (37). Studies have demonstrated that oxytocin has prosocial effects in adolescent and adult C58/J mice following a subchronic oxytocin regimen and that enhanced sociability was still present two weeks following treatment (38, 39). Acute oxytocin also significantly decreased abnormal repetitive behaviors in C58/J mice at doses that did not reduce general locomotion (38).

In the present study, we utilized the C58/J mouse model, with C57BL/6J (B6) serving as a comparison inbred strain, to evaluate the neurological mechanisms that contribute to sociability deficits and sex differences in ASD. C57BL/6J is a highly social strain that appears physically identical to C58/J and that shares genetic lineage with C58/J and thus is an appropriate control for C58/J (40). The 3-chamber social choice test was used to evaluate sociability in mice. Bulk RNA-Sequencing (bulk RNA-Seq) and single nucleus RNA-Sequencing (snRNA-Seq) of the amygdala, a brain region altered in individuals with ASD that connects specific neuroanatomical networks that regulate social function, was conducted to correlate transcriptomic profiles in this region to strain, sociability, sex, and the effects of oxytocin on behavioral phenotype (41-43). We interrogated DNA methylation differences in amygdala and analyzed mouse brain connectivity with Magnetic Resonance Histology (MRH). Our analysis of snRNA-Seq data determined that mature oligodendrocytes and microglia exhibit alterations in ASD risk gene expression, genes associated with myelination and microglia homeostatic control, gene regulatory networks, and cell differentiation that are associated with an ASD phenotype in C58/J mice that is dependent on sex and sociability. Bulk RNA-Seq analysis determined that oxytocin treatment had beneficial effects on myelin-related transcriptomic profiles in C58/J amygdala, while immune system-related biological processes that play a role in ASD differed between C58/J and C57BL/6J mice in amygdala. Differences in DNA methylation were also seen between C58/J and C57BL/6J amygdala. This work demonstrates the potential pathological roles of oligodendrocytes and microglia in ASD and provides insight into the mechanisms of oxytocin treatment.

## Methods

### Animals

C57BL/6J and C58/J mice were bred and tested for behavior at the University of North Carolina at Chapel Hill (UNC), with founder breeding pairs obtained from Jackson Laboratories (Bar Harbor, ME). Three separate sets of cohort groups were generated: 1) C57BL/6J and C58/J mice for a between-strain comparison of transcriptional and methylation profiles, 2) C58/J mice to investigate effects of oxytocin treatment on divergent social phenotypes (Supplemental Figure 1A-D); and 3) C58/J mice for MRH brain analysis. All mice were maintained at 20-23°C in groups of 2-4 in a specific pathogen-free room on 12 hr light and dark cycles with ad libitum access to food and water. All experimental procedures were conducted in compliance with an approved UNC IACUC protocol, and those set forth in the “Guide for the Care and Use of Laboratory Animals” as published by the National Research Council.

### Oxytocin Regimen

Oxytocin (Bachem, Torrance, CA) was dissolved in saline containing 0.002% glacial acetic acid. All injections were administered IP (intraperitoneal) in a volume of 10.0 ml/kg. Mice were given 4 injections of vehicle or oxytocin (1.0 mg/kg) over the course of post-natal weeks 6-7 with at least 48 hr between each injection. This regimen has previously been shown to have prosocial effects in mice (38, 39).

### Three-Chamber Choice Test

For mice used for the two-strain comparison or for MRI scans, sociability in the 3- chamber test was assessed one time, at age 7-8 weeks, as previously described (44). In the second set of mice, sociability was assessed 24 hr and 2 weeks following the final oxytocin or vehicle treatment, at ages 7-8 and 9-10 weeks. The test started with a 10 min habituation phase, with free exploration of the empty test box. Mice were then given a choice between an unfamiliar stranger mouse contained in a clear plexiglass cage in one side of the test box, or an empty plexiglass cage in the opposite side. Holes were drilled into the cages to allow olfactory investigation of the stranger mouse. Measures were taken of time spent in close proximity (within 5 cm) to the cage containing the stranger mouse or the empty cage by an automated image tracking system (Ethovision, Noldus Information Technology, Wageningen, the Netherlands). To study within-strain divergent social phenotypes, C58/J mice were divided, as evenly as possible, into four groups (male and female low social (ML, FL) and high social (MH,FH)), based on the percent of total proximity time each mouse spent in proximity to the stranger cage.

### Tissue Isolation and Processing

Perfusion, brain dissection, and tissue collection were performed one day after completion of the 3-chamber choice test in the first set of mice (C57BL/6J and C58J), and one day after the second 3-chamber test in the second set of mice treated with oxytocin or vehicle (C58/J). Mice were anesthetized with isoflurane prior to being perfused. Following perfusion, brains were rapidly removed from the skull and were placed into a chilled 1-mm stainless-steel brain matrix and divided into target regions using sterilized razor blades. The amygdala was collected within a 1.2-mm tissue punch. Tissue punches were placed in pre-chilled microcentrifuge tubes on dry ice and were stored at −80°C.

### Bulk RNA-Seq Library Preparation, Sequencing, Alignment, And Analysis

For bulk RNA-Seq analysis in the first set of mice, comparisons were made based on strain, sociability, and sex with C57BL/6J mice (males n = 4, females n = 4) serving as high-sociability controls, and C58/J mice separated into male and female low sociability (ML, FL, n = 4 each) and high sociability (MH, FH, n = 4 each) groups. In the second set of mice (all C58/J), comparisons were made based on oxytocin treatment, sociability, and sex with the following groups: ML Vehicle (n = 3), ML Oxytocin (n = 4), MH Vehicle (n = 5), MH Oxytocin (n = 4), FL Vehicle (n = 4), FL Oxytocin (n = 4), FH Vehicle (n = 5), and FH Oxytocin (n = 4). For each group, litter effects were controlled by taking only one mouse of each sex per sociability group (low or high) per litter, with maximum of 3 mice per litter.

DNA and RNA were extracted simultaneously from each sample using the All-Prep DNA/RNA Micro Kit (Qiagen, Hilden, Germany) according to the manufacturer’s instructions. Extracted DNA and RNA samples were quantified on a single-channel spectrophotometer to assess quality by 260/230 and 280/260 absorbance ratios and to determine their approximate yield. RNA samples were stored at −80°C and DNA was stored at 4°C. For library preparation, RNA samples were thawed on ice and concentrations were quantified using the Quant-iT™ RiboGreen™ RNA Assay Kit (ThermoFisher, Waltham, MA) and normalized to 10 ng/µL. RNA quality was assessed using high sensitivity RNA ScreenTape analysis run on a 4200 TapeStation (Agilent Technologies, Santa Clara, CA). Libraries were prepared using the TruSeq Stranded mRNA LP kit (Illumina, San Diego, CA) and indexed using the Illumina IDT-TruSeq RNA UD Idx kit. Quality control (QC) of prepared libraries was performed using Agilent high sensitivity D1000 ScreenTape analysis. Completed libraries were transferred to the Duke University Sequencing Core and sequenced on an Illumina NovaSeq 6000 S1 full flow cell. MultiQC reports (45) were generated prior to and after adapter trimming and phredbased read filtering using cutadapt. STAR (46) was used to align the reads to the GRCm38 mouse reference genome. The number of reads within each annotated mouse transcript were calculated using FeatureCount (47). PCAtools package version 2.2 was used to identify outlier samples and to determine factors impacting variation within the expression data. Differential expression analysis was performed using DESeq2 software (48). Genes with FDR below 0.01 and absolute fold change ≥ 0.5 were considered differentially expressed. Gene-set enrichment analysis (GSEA) was used for biological pathway analysis (49).

### Single Nucleus Preparation

High quality nuclei were isolated from 10-20 mg fresh frozen amygdala tissue pieces using 10x Genomics Chromium Nuclei Isolation Kit (10x Genomics, Pleasanton, CA). Tissue pieces were homogenized by grinding with a pestle and incubated in Lysis Buffer for 10 min on ice. Samples were then dissociated by pipetting, added to a Nuclei Isolation Column, and centrifuged at 16,000 x g for 20 sec at 4°C. The flow through was vortexed, centrifuged, and the nuclear pellet was then washed and centrifuged at 700 x g for 10 min at 4°C. Nuclear pellets were resuspended in Nuclei Wash and Resuspension buffer. Nuclei were counted and samples were adjusted to 1000 nuclei/μL using a Cellometer K2 cell counter (Nexcelom, Lawrence, MA).

### snRNA-Seq Library Preparation, Sequencing, And Analysis

snRNA-Seq libraries were constructed using 10x Genomics Chromium Single Cell 3’ Library & Gel Bead Kit v3 according to the manufacturer’s instructions. Nuclei from 10,000 cells were combined with reverse transcription (RT) reagents and loaded per channel on a 10x Genomics chromium controller with individually barcoded gel beads and oil to partition each sample into nanoliter-scale gel beads in emulsions (GEMs) within which the RT reaction occurs. GEMs were then broken to reveal full length cDNAs that were purified, amplified, and enzymatically fragmented before Illumina P5 and P7 sequences, sample indexes, and TruSeq Read 2 sequencing primers were added via end repair, A-tailing, adapter ligation, and PCR. cDNAs and final libraires were run on an Agilent 4200 TapeStation to check for quality and were quantified using KAPA Library Quantification Kit (Roche, Basel, Switzerland).

Libraries were transferred to the Duke University Sequencing Core and sequenced using paired end sequencing on an Illumina NovaSeq 6000 S1 full flow cell. Sample demultiplexing, barcode processing, and single-cell 3’ gene counting were performed using 10x Genomics Cell Ranger software. Reads were aligned to the GRCm38 mouse reference genome. The snRNA-Seq data (Cell Ranger result) contained 4 samples from amygdala (ML, MH, FL, FH). The resulting matrix files were used for subsequent bioinformatics analyses in Seurat (version 4.3.0) and R (version 4.1.1). Doublets were removed with the DoubletFinder package (version 2.0.3) and cells with less than 200 genes and more than 5% mitochondria gene count were excluded from analysis due to low quality. Data sets were normalized, logarithmically transformed, and subjected to Principal Component Analysis (PCA). Uniform manifold approximation and projection (UMAP) was utilized to visualize cell clustering. Manual annotation of cell clusters was done by identifying cell types through expression of canonical marker genes (50, 51). Differentially expressed genes were detected by default Wilcoxon rank sum test. GO enrichment analysis was performed using the clusterProfiler package (version 4.2.2) to identify enriched cellular pathways. The snRNA-Seq datasets were analyzed with scVelo to determine RNA Velocity (52, 53).

### DNA Methylation Analysis

Genomic DNA was isolated as described above. DNA (1 ug) was taken to the Duke University Center for Genomic and Computational Biology (GCB) and sheared to achieve 175 bp-long fragments using Covaris S220 (Covaris, Woburn, MA). DNA quality was assessed using Agilent 4200 TapeStation. Libraries were prepared using the Agilent SureSelect^XT^ Methyl-Seq Target Enrichment System. DNA fragments were bead-purified followed by end-repairing and A-tailing. After ligation of methylated adapters, the EZ-DNA Methylation Gold kit (Zymo Research, Irvine, CA) was used to perform bisulfite conversion. After post-bisulfite conversion cleanup, libraries were amplified using PCR and amplicons were purified with AMPure XP beads (Beckman Couter, Brea CA) and indexed with Agilent SureSelect^XT^ Index Set ILM. Quality control of libraries was done using Agilent high sensitivity D1000 ScreenTape and TapeStation analysis. Libraries were then sequenced on an Illumina NovaSeq S1 full flow-cell. Reads from the DNA methylation libraries were processed and mapped to GRCm38 using Bismark version 0.22.3 (54) and bowtie2 version 2.4.1 (55). CpG methylation values were extracted from the aligned bisulfite converted genomes using the Bismarck methylation extractor function. The R package Rnbeads version 2.8 (56) was used to quality assess the DNA methylation datasets and identify differential methylation at individual CpG resolution. Methylation coverage files generated by the Bismarck methylation extractor were annotated via the Rnbeads pipeline using the GRCm38 reference genome. Differentially methylated regions were determined using the R package dmrseq (57) and the BSmooth R package (58). CpGs were considered differentially methylated at an FDR (q score) > 0.1.

### SCENIC Analysis

The R packages SCENIC (version 1.1.2-01), RcisTarget (version 1.20.0), and AUCell (version 1.22.0) were used to identify transcription factors and cell states in snRNA-Seq data from C58/J amygdala (59). Raw UMI counts from Seurat served as input matrices for each sample in SCENIC. A gene filter was applied that kept genes with at least 6 UMI counts across all samples that were detected in at least 1% of the cells. GENIE3 (version 1.22.0) was used to identify potential transcription factor targets. The activity of each regulon was evaluated using AUCell which calculates the area under the recovery curve and integrates the expression ranks across all genes in a regulon. Cells were clustered according to gene regulatory network or regulon activity.

### Cell-To-Cell Communication Analysis

The R package CellChat (version 1.6.0) was used to analyze intercellular communications within the snRNA-Seq datasets from C58/J amygdala (60). CellChat is a public database of ligands, receptors, cofactors, and their interactions. The CellChat R toolkit and a Web-based “CellChat Explorer” (http://www.cellchat.org/) were used to identify intercellular communications and help construct cell-cell communication atlases. For the cell-interaction analyses, the expression levels were calculated relative to the total read mapping to the same set of coding genes in all transcriptomes. The expression values were averaged within each single-cell cluster or cell sample.

### Magnetic Resonance Histology

C58/J mice underwent transcardial perfusions with a mixture of 10% buffered formalin and Prohance (Gadoteridol) to reduce the spin lattice relaxation time and enhance the signal for MRH (61). Images were acquired on a 7 T horizontal bore magnet with Agilent Direct Drive console and Resonance Research high performance gradients with peak gradients of 2000 mT/m. The head of the perfused specimen with the brain in the cranial vault was placed in a 12 mm diameter solenoid radiofrequency coil. Diffusion tensor MR images were acquired using a Stesjkal Tanner spin echo imaging sequence with TR/TE= 100/15.8 ms with diffusion weighting (bvalue) of 3000 s/mm^2^. Forty-six diffusion weighted images were acquired using gradient vectors equally distributed on the unit sphere. Baseline (b0) images were acquired after every tenth volume. The acquisitions were accelerated using compressed sensing with a compression factor of 8 resulting in isotropic resolution of 35 μm with acquisition time of 22 hrs. Volumes were registered together, denoised and processed using a series of imaging pipelines described fully in (62). Image volumes were registered to a standardized atlas with a label set consistent with the ABA CCFv3 (62). Statistical comparisons between groups (ML, MH, FL, FH) were performed using Omni-MANOVA analysis (63). Composite images were created for each group using the QSDR method (64).

## Results

### C58/J mice display divergent levels of sociability and differentially expressed genes associated with ASD compared to C57BL/6J mice

Our study replicated previous findings of high and low sociability phenotypes within the isogenetic C58/J inbred strain (44). Notably, the high sociability C58/J mice spent similar amounts of time in proximity to the stranger mouse as the C57BL/6J controls, while the low sociability groups had significantly lower duration than both the high-sociability C58/J mice and C57BL/6J (Figure 1A, see also Supplemental Figure 1). Bulk RNA-Seq differential gene expression analysis was conducted in the amygdala to gain insight into the molecular mechanisms that contribute to strain-level differences in sociability between C58/J and C57BL/6J mice. With an adjusted p-value < 0.05 and log2-FoldChange of 2 and −2, the bulk RNA-Seq results identified 64 upregulated and 63 downregulated genes in C58/J amygdala (Figure 1B).

**Figure 1.**
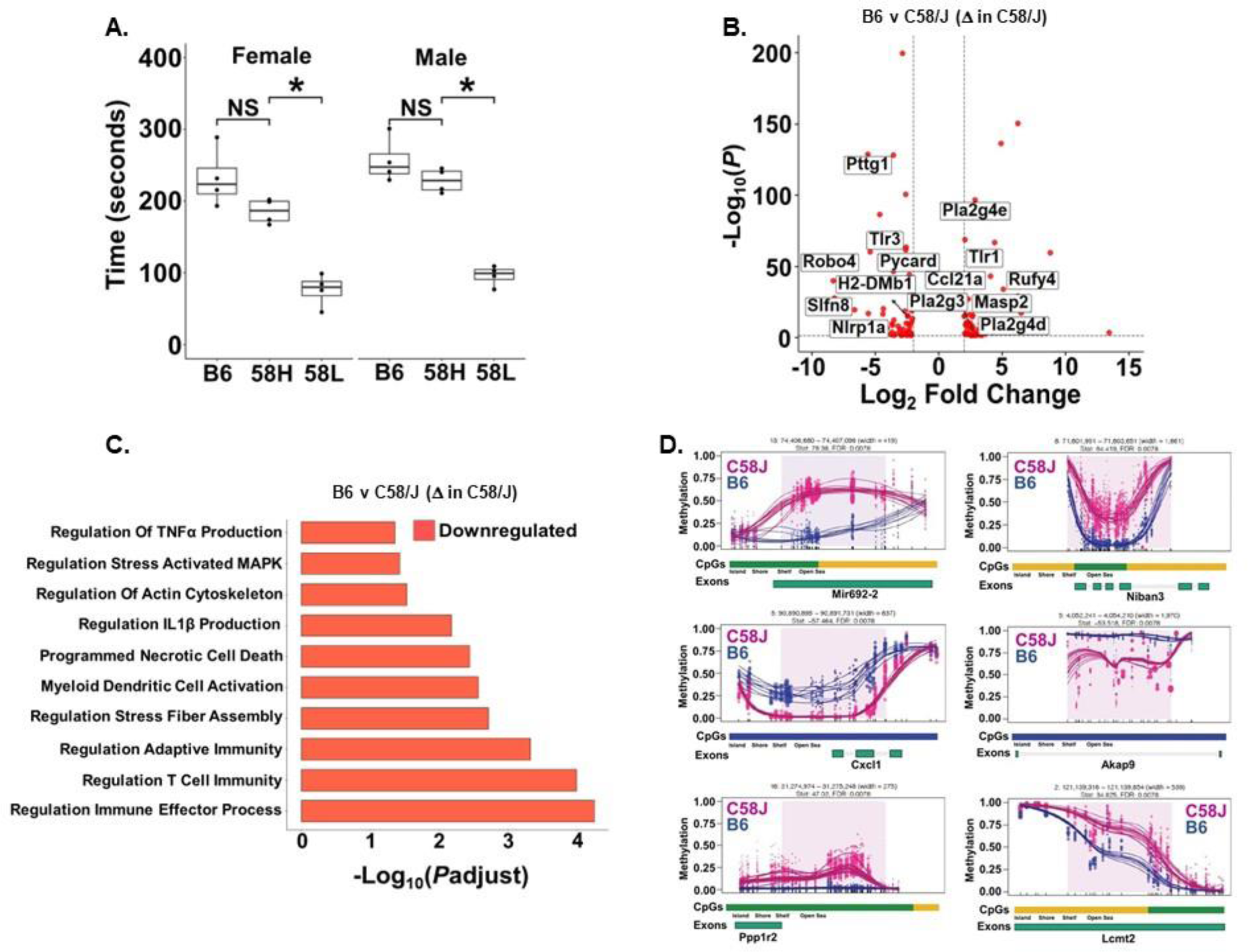
C58/J mice exhibit reduced sociability and downregulation of immune-related pathways in amygdala compared to C57BL/6J mice. (A) Comparison of social preference based on time spent in close proximity to a stranger mouse during the 3-chamber social choice test in male (n = 4 per group) and female (n = 4 per group) C57BL/6J (B6), C58/J high social (58H), and C58/J low social (58L) mice. Preference for social interaction was assessed at postnatal weeks 7-8. Results reported as mean ± SEM, *p<0.05 using Mann-Whitney U test. (B) Volcano plot of differentially expressed genes obtained from bulk RNA-Seq analysis of amygdala from C58/J and C57BL/6J mice (n = 4 mice per group). Genes with an adjusted *p*-value < 0.05 and Log2 Fold Change of > 2 and < −2 were differentially expressed. 64 genes were upregulated and 63 genes were downregulated. (C) GO enrichment analysis of bulk RNA-Seq data from C58/J and C57BL/6J amygdala. Pathways were downregulated in C58/J mice. (D) Significant (*q*-value < 0.05) hypermethylated (*Mir692-2*, *Niban3*, *Ppp1r2*, *Lcmt2*) and hypomethylated (*Cxcl1*, *Akap9*) regions identified in C58/J compared to C57BL/6J mice (n = 4 mice per group). Lines represent individual smoothed methylation level estimates for C58/J (C58J, red) or C57BL/6J (B6, blue). Dots represent methylation level estimates of an individual CpG in a single sample, and dot size is representative of coverage. CpG and genic annotation tracks are shown below each plot.

Many of the differentially expressed genes (DEGs) identified were associated with immune function and evidence highlights a link between immune dysfunction and ASD (65). For instance, elevated levels of the enzyme Phospholipase A2 (Pla2) are associated with ASD, while *Pla2* genes have been shown to be differentially expressed in ASD (66, 67). *Pla2g4d* and *Pla2g4e* were upregulated in C58/J amygdala, while *Pla2g3* was downregulated in comparison to C57BL/6J (Figure 1B). Studies also suggest that innate-immunity-related pathways play an important role in autism and have identified several genes in these pathways that are differentially expressed in ASD (68). Multiple DEGs in C58/J amygdala encode proteins that are part of innate-immunity-related pathways including *Treml2* (myeloid cell innate immunity), *Pycard* (Nod-like receptor signaling), *Tlr3*/*Tlr1* (Toll-like receptor signaling), and *Ccl21a* (Chemokine signaling) (Figure 1B).

Like innate immunity, neuronal migration plays an important role in the developing brain and is often impaired in ASD. Recent studies have shown that neuronal migration is reduced in mouse brain when members of the brain complement pathway, such as Masp1 and Masp2, are decreased and our findings show *Masp2* gene expression is increased in C58/J amygdala (69) (Figure 1B). Genetic analyses indicate the axon guidance molecule *Robo4* may play a role in ASD by affecting serotonin signaling or neurodevelopment and *Robo4* was overtly downregulated in C58/J amygdala (70) (Figure 1B). Finally, autophagy is a degradation mechanism that helps maintain neuronal homeostasis, survival, and plays a role in the progression of neuronal disease. The *Rufy4* gene that encodes a protein responsible for macroautophagy was highly upregulated in C58/J amygdala (71) (Figure 1B). The analysis of enriched Gene Ontology (GO) signaling pathways revealed many of the downregulated genes were enriched in immune-related biological processes such as regulation of TNFα and IL1β production, regulation of T Cell immunity, programmed necrotic cell death, regulation of adaptive immunity, and myeloid dendritic cell activation (Figure 1C).

In addition to alterations in gene expression and signaling pathways, epigenetic mechanisms may shed light on what distinguishes C58/J mice from C57BL/6J mice. Differential methylation analysis conducted using DMRseq between C58/J and C57BL/6J identified 248 differentially methylated regions (DMRs) in amygdala (100 hypomethylated, 148 hypermethylated). The DMRs in amygdala overlapped with 70 annotated genes. The top hypermethylated genes in C58/J compared to C57BL/6J amygdala overlapped with the following genes at *q*-value = 0.5 x 10^-2^: the micro-RNA *Mir692-2*, Apoptosis Regulator 3 (*Niban3*), Protein Phosphatase 1 Regulatory Inhibitor Subunit 2 (*Ppp1r2*), and Leucine Carboxyl Methyltransferase 2 (*Lcmt2*) (Figure 1D). Two of the top hypomethylated genes in C58/J relative to C57BL/6J amygdala were C-X-C Motif Chemokine Ligand 1 (*Cxcl1*) and A-Kinase Anchoring Protein 9 (*Akap9*) (Figure 1D). Several members of the A-kinase anchor protein (Akap) family, including Akap9, are functionally and genetically linked to ASD, while studies have shown levels of the immunological marker Cxcl1 were significantly elevated in ASD patients (72, 73).

In addition to attempting to uncover the molecular mechanisms that differentiate C58/J mice from C57BL/6J mice, we utilized MRH to examine variations of brain architecture in C58/J mice as a function of sex and sociability (Supplemental Figure 2). MRH analysis revealed no differences in whole-brain connectomes or in regional brain volumes between ML, FL, MH, and FH mice suggesting that divergent sociability could not be attributed to overt neuroanatomical or connectivity differences.

### C58/J mouse amygdala consists of 29 cell types with distinct ASD-like transcriptomic signatures that were dependent on sex and sociability

Mice were sacrificed and amygdala were collected for bulk RNA-Seq and snRNA-Seq analysis (Figure 2A). A quality-controlled single cell atlas of C58/J amygdala included 31,447 cells from ML, MH, FL, and FH mice. Unsupervised Seurat analysis revealed 29 distinct UMAP clusters (Figure 2B), which were annotated using canonical markers of the mouse cortex from previous studies (Table 1) (50, 51). These 29 cell populations could be simplified into 3 main cell types: glutamatergic (*Slc17a7*/*Slc17a6*, *Celf2*, *Arpp21*, *Pcsk2*, *Ptprd*, *Ano3*) (Supplemental Figure 3A-B), GABAergic (*Adarb2*+/*Adarb2*-, *Gad1*, *Gad2*, *Grip1*, *Dlx6os1*) (Supplemental Figure 3C-D), and non-neuronal (Supplemental Figure 3E). Based on the enriched gene features, the four C58/J groups showed notable variability of cell type proportions. For instance, the ML data contained elevated numbers of L2/3 IT CTX cells (40.3%, 2981 cells), while MH had a greater number of Meis2-Penk-GABA cells (13.5%, 963 cells) and the FH had increased L5 IT CTX Rspo1 cells (12.9%, 936 cells) (Supplemental Figure 4A-B). In addition, the MH and FL mice had a greater number of astrocytes (9.3%, 699 cells; 8.6%, 820 cells), microglia (5.6%, 400 cells; 5.0%, 475 cells) and mature oligodendrocytes (6.3%, 450 cells; 7.1%, 675 cells) compared to the ML (astrocytes 2.75%, 201 cells; mature oligodendrocytes 0.66%, 49 cells; microglia 1.0%, 74 cells) and FH (astrocytes 5.1%, 368 cells; mature oligodendrocytes 1.0%, 74 cells; microglia 1.2%, 89 cells) (Supplemental Figure 4A).

**Figure 2.**
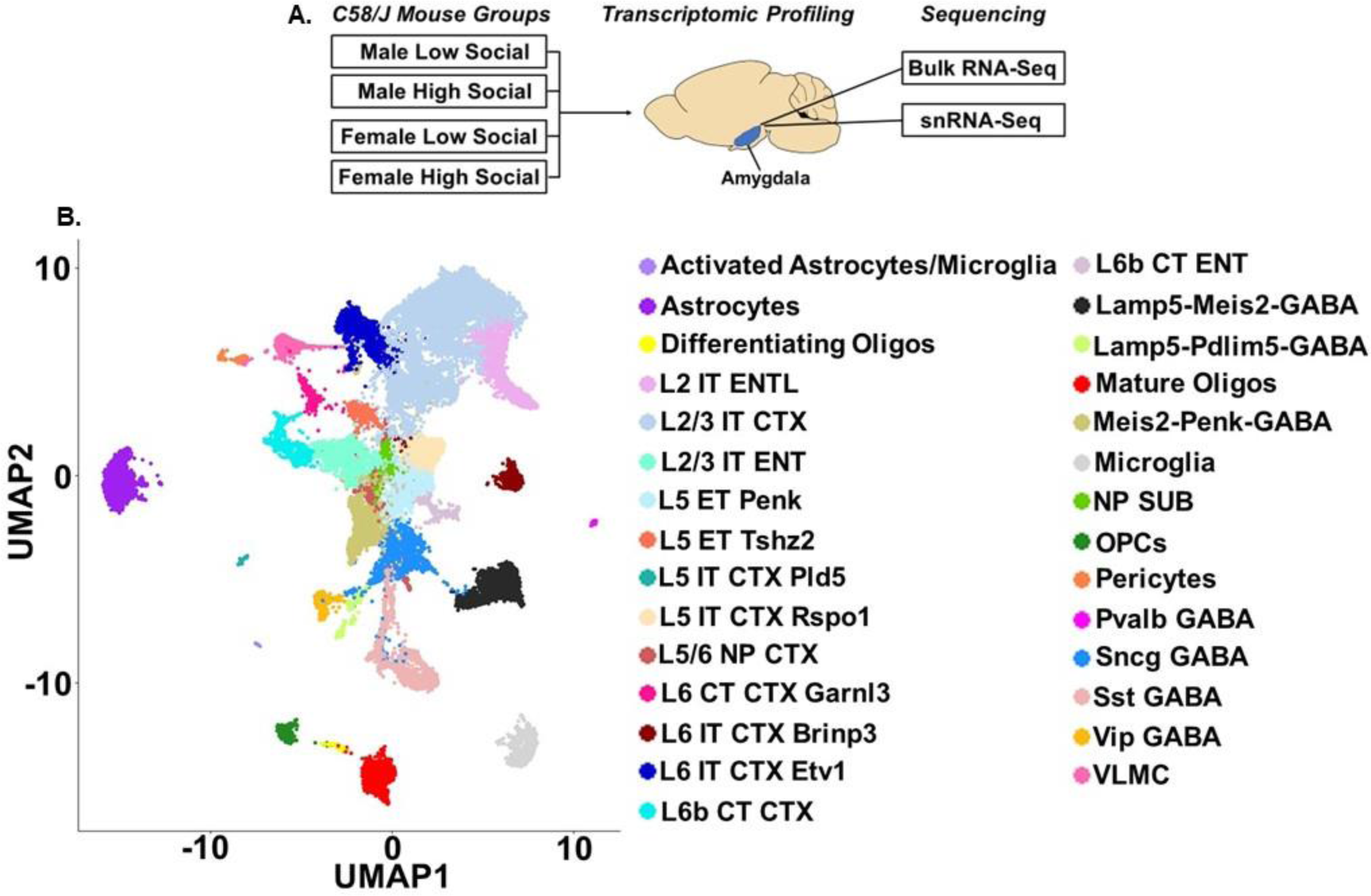
Clustering of 31,447 cells identified 29 cell types in C58/J amygdala. (A) Amygdala was dissected from brains taken from C58/J Male Low Social (ML), Male High Social (MH), Female Low Social (FL), and Female High Social (FH) mice (n = 1 mouse per group) and was used for bulk RNA-Sequencing (Bulk RNA-Seq) and single nucleus RNA-Sequencing (snRNA-Seq). (B) UMAP plot of 31,447 cells and the 29 distinct cell clusters that were obtained from snRNA-Seq.

**Table 1.**
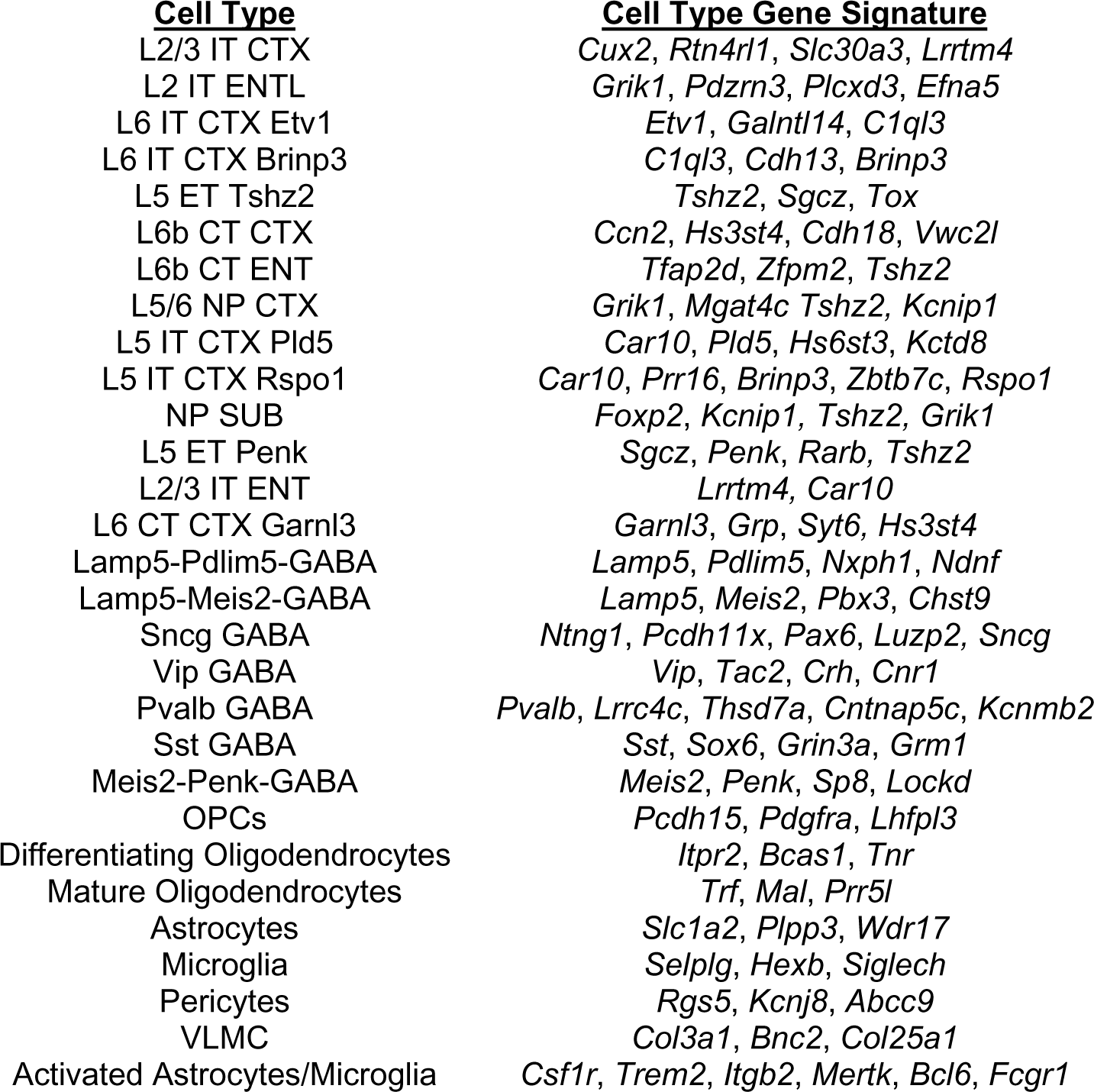
Cell Types and their gene signatures in C58/J amygdala.

Next, we stratified our snRNA-Seq and bulk RNA-Seq data based on sex (ML vs. FL, MH vs. FH) and sociability (ML vs. MH, FL vs. FH) given the core ASD epidemiological features of social dysfunction and a greater frequency in males. Bulk RNA-Seq comparison between MH vs. FH showed the greatest differences in gene expression (239 genes) compared to the other groups (ML vs. FL 10 genes, ML vs. MH 45 genes, FL vs. FH 12 genes) (Supplemental Figures 5&6). These 239 genes included seven high confidence (HC) and strong candidate (SC) ASD risk genes from the Simons Foundation Autism Research Initiative (SFARI) ASD risk gene database (Supplemental Figure 6B). Pathway analysis of the bulk RNA-Seq DEGs revealed that many of the top enriched Gene Ontology (GO) terms in the MH vs. FH comparison could be functionally related to ASD pathogenesis, including synapse organization, learning, protein demethylation, axonogenesis, and synaptic plasticity (Figure 3B).

**Figure 3.**
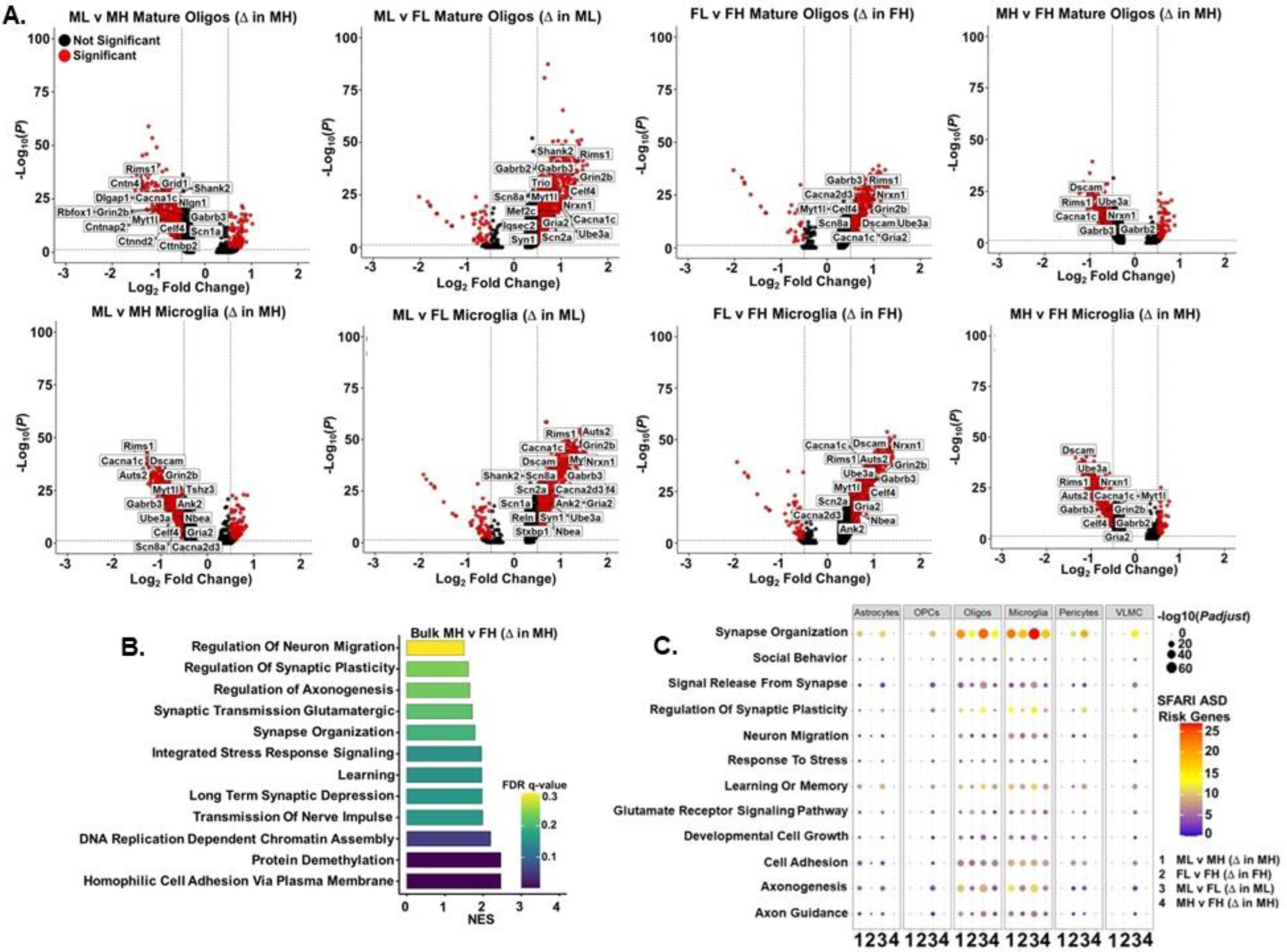
Mature oligodendrocytes and microglia from Male Low Social and Female High Social C58/J amygdala demonstrate the greatest number of differentially expressed SFARI ASD risk genes. (A) Volcano plots showing differentially expressed genes (DEGs) obtained in microglia and mature oligodendrocytes when C58/J mouse groups (n = 1 mouse per group) were compared based on sex and sociability. Specific high confidence (HC) SFARI ASD risk DEGs are labeled in the plots. (B) GO Biological Process (GO BP) enrichment analysis of ASD-related pathways in MH amygdala compared to FH amygdala obtained by bulk RNA-Seq analysis and GSEA (n = 5 mice per group). Positive normalized enrichment score (NES) indicates pathways are upregulated in MH. (C) Dot plots of GO BP analysis of ASD-related pathways for six non-neuronal cell types when C58/J mouse groups were compared based on sex and sociability. Dot color indicates number of HC and SC SFARI ASD risk genes. Dot sizes are proportional to – log10(*P*adjust).

Analysis of the snRNA-Seq data revealed DEGs and SFARI risk genes within ML and FH cell types (Supplemental Figures 5&6). Interestingly, mature oligodendrocytes and microglia contained the greatest number of DEGs across the four comparisons (Supplemental Figures 5&6). Compared to the FL and MH groups, ML and FH mature oligodendrocytes and microglia showed the greatest differences in gene expression (Figure 3A). GO term analysis of the snRNA-Seq dataset revealed that many ASD-related pathways were enriched specifically in microglia and mature oligodendrocytes with the greatest number of DEGs and SFARI risk genes occurring in the ML group when compared to MH (sociability) and FL (sex) (Figure 3C).

### Dysregulation of myelin-related gene expression and myelin-related pathways in mature oligodendrocytes from ML and FH C58/J amygdala

Next, we focused on the interactions between sex and sociability on oligodendrocyte expression given the differences of mature oligodendrocyte numbers in ML and FH together with the differential expression of ASD risk genes in this cell type. MH and FL oligodendrocytes were enriched in myelin-related genes, such as *Aspa*, *Mag*, *Mog*, and *Plp1,* which are characteristic of mature differentiated myelinating oligodendrocytes (Figure 4A) (74). GO pathway analysis showed enrichment in biological processes related to myelination, including neuron/axon ensheathment, fatty acid/ATP biosynthesis and myelination, and oligodendrocyte differentiation (Figure 4B). In contrast, ML and FH oligodendrocytes had elevated expression of genes like *Ptprz1*, *Tnr*, *Dscam*, *Sox5*, and *Sox6* that indicate an OPC-like phenotype (Figure 4A). Bulk RNA-Seq analysis by sex and sociability corroborated our findings related to myelination. To further investigate expression profiles associated with sociability, analyses were carried out for a separate set of C58/J mice given a subchronic regimen with either oxytocin or vehicle. Interestingly, FH mice treated with oxytocin (FHOXY) exhibited upregulation of genes involved in myelin synthesis (*Trf*, *Aspa*, *Plp1*, *Mog*, *Mal*, *Mobp*, *Myrf*, *Cnp*, *Mag*) in amygdala compared to vehicle-treated FH (FHC) mice (Figure 4C). Signaling pathways related to myelination were also upregulated in amygdala from FHOXY compared to FHV, as well as in MH amygdala compared to FH amygdala (Figure 4D).

**Figure 4.**
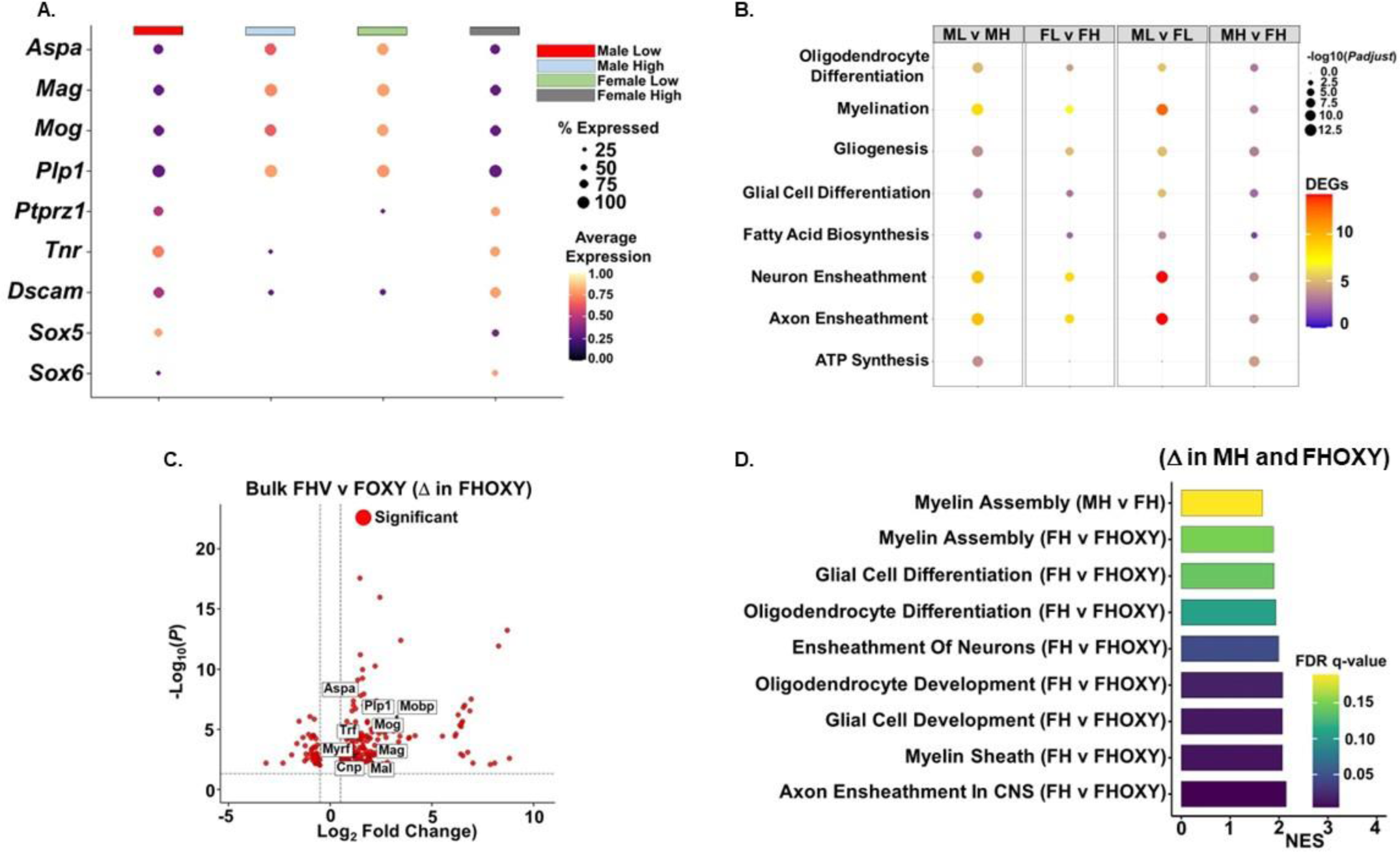
Dysregulation of myelinating gene expression and myelinating pathways in mature oligodendrocytes from Male Low Social and Female High Social C58/J amygdala. (A) Dot plot of genes in mature oligodendrocytes from representative groups of C58/J amygdala (n = 1 mouse per group) that are markers for mature oligodendrocytes (*Aspa*, *Mog*, *Mag*, *Plp1*) and OPCs (*Ptprz1*, *Tnr*, *Dscam*, *Sox5*, *Sox6*). Color intensity implies level of gene expression and dot size indicates the percentage of each cluster expressing the gene. (B) Dot plots of Gene Ontology Biological Process (GO BP) analysis of myelination-related pathways in mature oligodendrocytes from representative groups of C58/J amygdala that were compared based on sex and sociability. Dot color indicates number of DEGs in the pathway. Dot sizes are proportional to –log10(*P*adjust). (C) Volcano plot showing differentially expressed genes (DEGs) obtained by bulk RNA-Seq in Female High Social amygdala vehicle (FHV) and oxytocin-treated (FHOXY) groups (n = 4-5 mice per group). Specific myelinating genes are labeled in the plot. Genes are differentially expressed in the FHOXY group. (D) GO BP enrichment analysis of pathways involved in myelination in Male High Social (MH) amygdala compared to Female High Social (FH) amygdala and FHV amygdala compared to FHOXY amygdala obtained by bulk RNA-Seq analysis and GSEA (n = 4-5 mice per group). Positive normalized enrichment score (NES) indicates pathways are upregulated in MH and FHOXY groups.

Cell-cell communication plays an important role in coordinating processes such as play a more important signaling role in mature oligodendrocytes and OPCs in ML compared to FL and MH (Supplemental Figure 7). Like ML, pericytes in FH signal to mature oligodendrocytes and OPCs using Wnt, Bmp, laminin and Vtn signaling, but they also signal via Spp1 which is pro-inflammatory (Supplemental Figure 7). and FH which have fewer mature oligodendrocytes (Figure 5A-D) (76). BMPs are growth factors that are members of the transforming growth factor beta (TGF-β) superfamily and inhibition of BMP signaling has been shown to induce oligodendrogenesis and remyelination (77). ML and FH pericytes signaled through multiple Bmp ligand-receptor pairs to OPCs and mature oligodendrocytes, while Bmp ligand-receptor pairs were not prominent in MH and FL (Figure 5A-D). The timing of oligodendrocyte differentiation and prominent in MH and were not present in FL or FH (Figure 5A-D). Studies suggest moderate Wnt signaling plays a role in oligodendrocyte differentiation (79). In ML and FH, pericytes used Wnt4-based signaling to communicate with both mature oligodendrocytes and OPCs while MH and FL did not demonstrate this (Figure 5A,D). Finally, laminin regulates oligodendrocyte survival, migration, proliferation, differentiation, and myelination (80). Compared to ML, more cell types in MH utilized Lama4-based ligand-receptor pairs to signal to OPCs and mature oligodendrocytes (Figure 5A-B).

**Figure 5.**
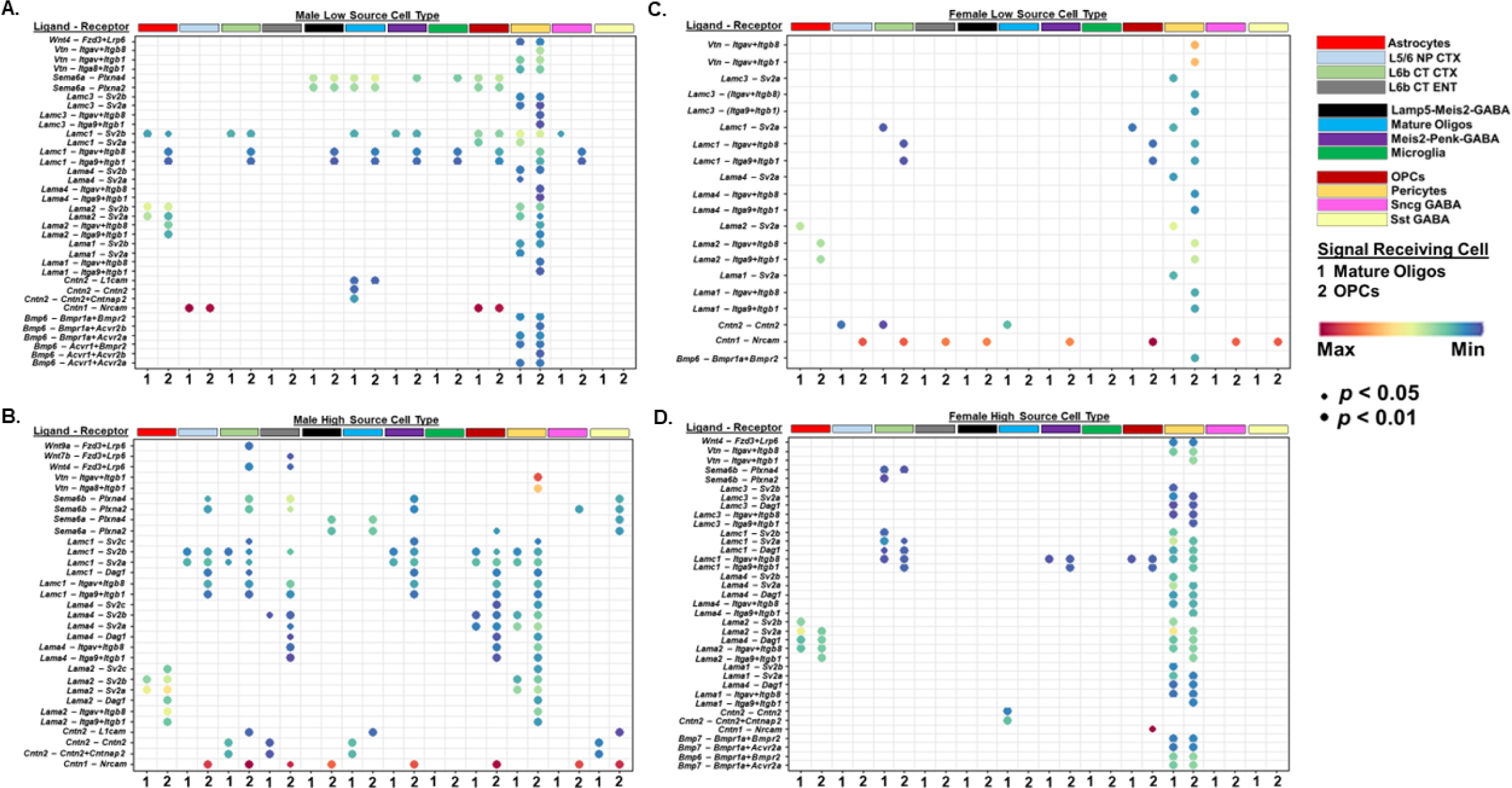
Ligand-Receptor pairs that contribute to signals originating from specific cell types that are received by mature oligodendrocytes and OPCs in C58/J amygdala. (A-D) Dot plots obtained via CellChat showing unidirectional cell-cell ligand receptor interactions. Ligands originate from the indicated 12 source cell types and interact with receptors in OPCs (2) and mature oligodendrocytes (1) from representative groups of C58/J amygdala (n = 1 mouse per group). Color implies expression magnitude and dot size indicates specificity of the interaction.

### Homeostatic gene expression and cell-cell communication are altered in microglia from ML and FH C58/J amygdala

Like mature oligodendrocytes, reductions in cell density and upregulation of ASD risk genes were observed in ML and FH microglia. Microglia in ML and FH exhibited reduced levels of homeostatic genes such as *C1qa, C1qb, Ctss, Cx3cr1, Entpd1, Gpr34, Selplg, Siglech, Slc2a5, Sparc, and Trem2* (Figure 6A) (81). GO pathway analysis revealed oxidative phosphorylation (OXPHOS), aerobic respiration, regulation of the immune system, and synaptic pruning were elevated in MH microglia compared to ML microglia suggesting microglia were more quiescent in MH compared to ML (Figure 6B). Synaptic pruning and several immune-related pathways were also elevated in FL microglia compared to ML microglia (Figure 6B). Interestingly, pathways related to the neuroinflammatory response and regulation of the immune system were elevated in FH microglia compared to FL microglia (Figure 6B). Pathway analysis of bulk RNA-Seq data supported our snRNA-Seq data in that differences were observed in immune system-related pathways and OXPHOS between MH and FH amygdala (Figure 6C).

**Figure 6.**
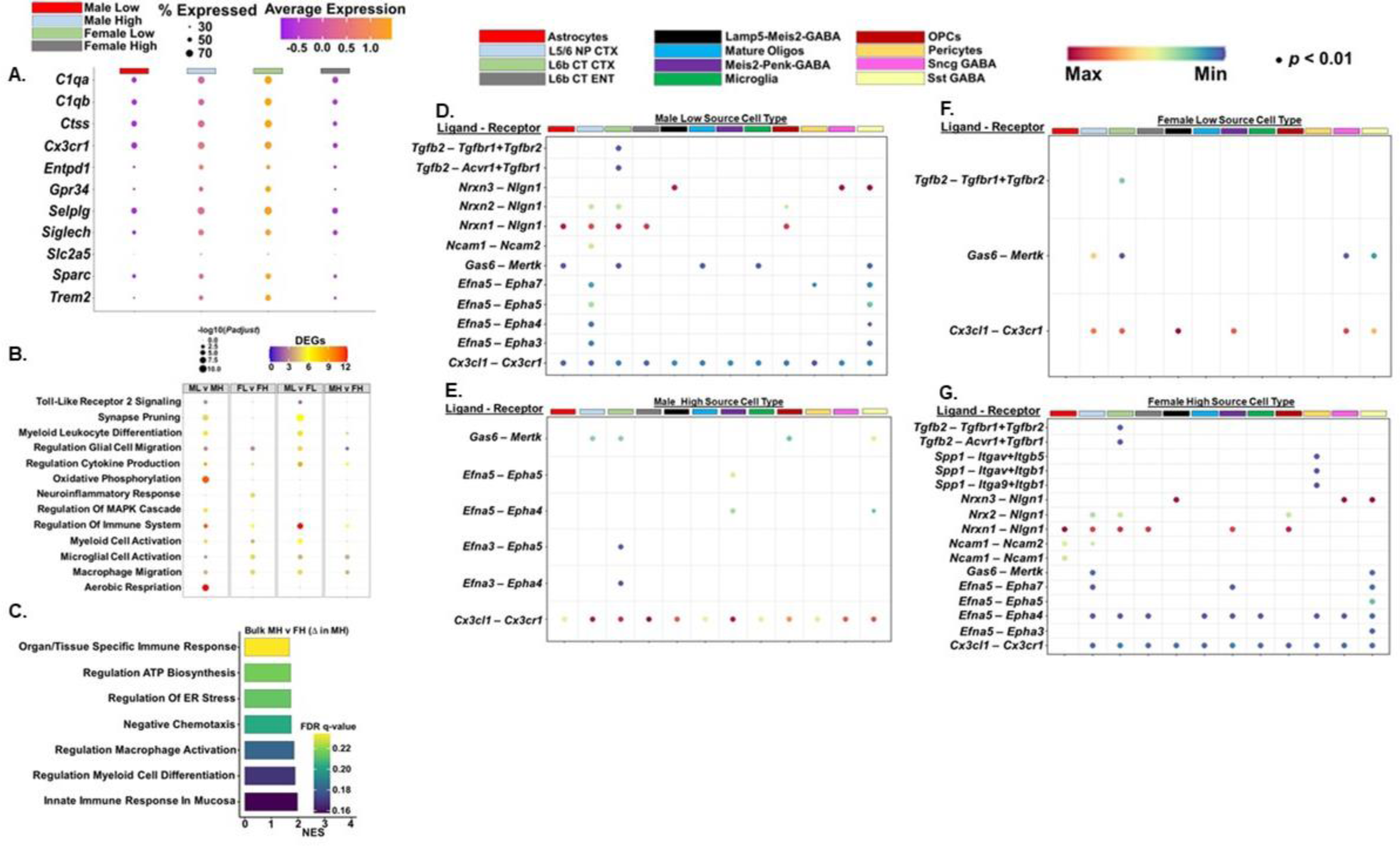
Homeostatic gene expression and cell-cell communication are altered in microglia from Male Low Social and Female High Social C58/J amygdala. (A) Dot plot of genes in microglia from representative groups of C58/J amygdala (n = 1 mouse per group) that are markers for microglial homeostasis. Color intensity implies level of gene expression and dot size indicates the percentage of each cluster expressing the gene. (B) Dot plots of GO Biological Process (GO BP) analysis of immune/microglial-related pathways in microglia from representative groups of C58/J amygdala that were compared based on sex and sociability (n = 1 mouse per group). Dot color indicates number of differentially expressed genes (DEGs) in the pathway. Dot sizes are proportional to – log10(*P*adjust). (C) GO BP enrichment analysis of immune-related pathways in Male High Social (MH) amygdala compared to Female High (FH) social amygdala obtained by bulk RNA-Seq analysis and GSEA (n = 5 mice per group). Positive normalized enrichment score (NES) indicates pathways are upregulated in the MH group. (D-G) Dot plots obtained via CellChat showing unidirectional cell-cell signaling originating from the indicated 12 source cell types in C58/J amygdala that release ligands that interact with receptors on microglia (n = 1 mouse per group). Color implies expression magnitude and dot size indicates specificity of the interaction.

We utilized CellChat to analyze the pathways used by different cell types to signal to microglia (Figure 6D-G). Neuronal cell adhesion molecules (Ncams) are cell surface proteins that function in neurodevelopment and studies suggest play a role in ASD-related neuroinflammation (82). Our results revealed different cell types utilized Ncam1-Ncam1 and Ncam1-Ncam2 ligand receptor pairs to signal to microglia in FH and ML (Figure 6D,G). Osteopontin (Spp1) is a cytokine involved in several physiological and pathophysiological processes including inflammation and macrophage activation (83, 84). In FH, pericytes utilized Spp1-(Itgav+Itgb5), Spp1-(Itgav-Itgb1), and Spp1-(Itga9+Itgb1) ligand-receptor pairs to signal to microglia (Figure 6G). In ML and FH, several cell types utilized Nrxn1-Nlgn1 ligand-receptor pairs to signal to microglia (Figure 6D,G). Neurexins (Nrxn) play a role in synapse formation and synaptic transmission and these findings suggest neural connectivity issues may be present in ML and FH microglia. Finally, Ephrin receptors and their ligands play a role in cell differentiation, positioning, and migration during nervous system development (85). Many ML and FH cell types utilized Efna5-Epha7, Efna5-Epha5, Efna5-Epha4, and Efna5-Epha3 ligand receptor pairs to signal to microglia compared to FL and MH (Figure 6D-G). In addition to analyzing signaling pathways utilized by different cell types to communicate with microglia, we also analyzed the pathways used by microglia to signal to other cell types (Supplemental Figure 8A-D). Ephrin, neurexin, and Ncam outgoing signaling from microglia to other cell types was much more prominent in ML and FH.

### Mature oligodendrocytes and microglial cells have distinct regulon activities that are driven by differences in sex and sociability in C58/J amygdala

The maintenance of cell identity involves the coordinated action of transcription factors (TFs) which regulate gene regulatory networks (GRNs) and control gene expression in cells. We utilized SCENIC to computationally reconstruct GRNs in mature oligodendrocytes and microglia separately based on single nuclei RNA expression data from C58/J amygdala to identify the key TFs that regulate these two cell types. SCENIC is comprised of three steps that include co-expression analysis, target gene motif enrichment analysis, and assessment of regulon activity (59). SCENIC lists regulons that represent a TF and significantly enriched target genes, as well as regulon activity scores in each cell. We were particularly interested in determining whether these two cell types have different gene regulatory circuitries across our four groups of C58/J amygdala. Our analysis indicates that GRN differences in C58/J amygdala mature oligodendrocytes and microglia are driven by differences in mouse sociability and sex. Following construction of the GRNs, regulon activities in the dataset were binarized into “on/off” for easier interpretation. Our network analysis revealed *Sox10* was an active regulon in FH mature oligodendrocytes (Figure 7A). Members of the Sox family of TFs (e.g., *Sox10*, *Sox8*) are key regulators of OPC specification, proliferation, migration, terminal differentiation into mature myelinating oligodendrocytes, and myelin maintenance (86). *Sox10* expression was detected in FH mature oligodendrocytes (Figure 7B) where many *Sox10*-regulated genes were upregulated compared to both MH and FL mature oligodendrocytes many of which were genes strongly associated with OPC identity such as *Pcdh15*, *Opcml*, *Ptprz1*, *Lhfpl3*, *Dscam* and *Tnr* (Figure 7C). In FL mature oligodendrocytes, *Sox10* was also an essential regulator (Figure 7A). *Sox10* was expressed in FL mature oligodendrocytes (Figure 7B) and the *Sox10* motif is shown in Figure 7E. Many *Sox10*-regulated genes were upregulated in FL mature oligodendrocytes compared to both FH and ML mature oligodendrocytes including genes that are associated with mature myelinating oligodendrocytes such as *Plp1*, *Mag*, *Mog*, *Mal*, *Cldn11*, and *Trf* (Figure 7D). Our analysis also identified 21 regulons that were active in FL microglia many of which regulated the expression of genes that were differentially expressed in FL microglia compared to FH and ML microglia including the *Irf8* and *Mafb* regulons (Figure 7A). Runx1 is a member of the Runx protein family and promotes microglia maturation as well as maturation of non-myelinating axon-wrapping glial cells in olfactory nerves (87, 88). *Runx1* was expressed in FL microglia and is predicted to regulate the expression of a number of differentially expressed genes in FL microglia compared to ML and FH microglia including the microglial homeostatic genes *Gpr34*, *Serinc3*, *Sparc*, *Cx3cr1*, and *Lgmn* (Figure 7F- G). The *Runx1* motif is shown in Figure 7I. *Runx1* was also expressed in MH microglia (Figure 7F) and was an active regulon that regulated the expression of many genes that were upregulated in MH microglia compared to FH and ML microglia including *Gpr34* and *Cx3cr1* (Figure 7H), while *Mafb* regulated the expression of many microglial homeostatic genes that were upregulated in MH microglia including *P2ry12* and *Gpr34*. Finally, *Sox8* was an active regulon in MH mature oligodendrocytes (Figure 7A). *Sox8* was expressed in MH mature oligodendrocytes (Figure 7J) and regulated the expression of many genes that were upregulated in MH mature oligodendrocytes compared to ML and FH mature oligodendrocytes including *Plp1*, *Cnp*, *Mal*, *Mog*, and *Mag* which are all associated with mature myelinating oligodendrocytes (Figure 7K). The *Sox8* motif is shown in Figure 7L.

**Figure 7.**
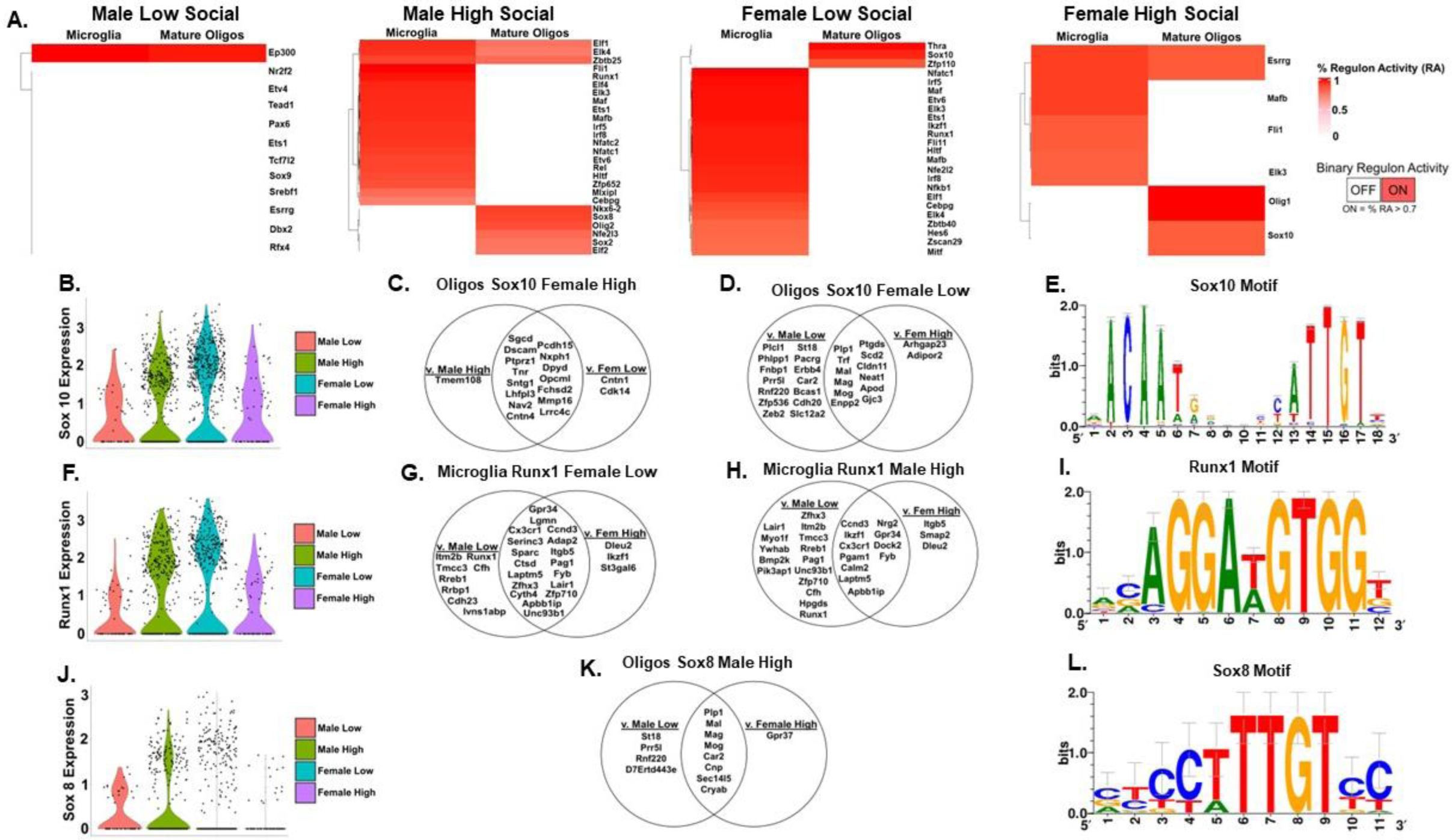
Mature oligodendrocytes and microglial cells have distinct regulon activities in C58/J amygdala that are dependent on mouse sex and sociability. (A) Binary regulon activity generated using SCENIC for microglia and mature oligodendrocyte cell clusters from amygdala of Male Low Social, Male High Social, Female Low Social, and Female High Social C58/J mice (n = 1 mouse per group). To determine active regulons in each cluster, an AUC threshold of 0.7 was used. Red blocks in the heatmap represent regulons whose activity met that threshold and are “on”, while white blocks represent regulons with activity below this threshold and are “off”. (B) Violin plot of Sox10 expression in mature oligodendrocytes from representative C58/J amygdala samples. (C) Venn diagram of differentially expressed genes (DEGs) in Female High Social mature oligodendrocytes compared to Male High Social and Female Low Social mature oligodendrocytes that are regulated by the Sox10 transcription factor. (D) Venn diagram of DEGs in Female Low Social mature oligodendrocytes compared to Male Low Social and Female High Social mature oligodendrocytes that are regulated by the Sox10 transcription factor. (E) Sox10 motif. (F) Violin plot of Runx1 expression in microglia from representative C58/J amygdala samples. (G) Venn diagram of DEGs in Female Low Social microglia compared to Male Low Social and Female High Social microglia that are regulated by the Runx1 transcription factor. (H) Venn diagram of DEGs in Male High Social microglia compared to Male Low Social and Female High Social microglia that are regulated by the Runx1 transcription factor. (I) Runx1 motif. (J) Violin plot of Sox8 expression in mature oligodendrocytes from representative C58/J amygdala samples. (K) Venn diagram of DEGs in Male High Social mature oligodendrocytes compared to Male Low Social and Female High Social mature oligodendrocytes that are regulated by the Sox8 transcription factor. (L) Sox8 motif.

### RNA velocity analysis reveals impaired differentiation of microglia and oligodendrocytes in ML and FH amygdala.

We performed RNA velocity analysis to investigate the transition states of oligodendrocytes and microglia in C58/J amygdala (52, 53). Latent time is a measure that demonstrated that oligodendrocyte cells are ordered along a differentiation trajectory with OPCs as the initial state followed by pODs and then mature oligodendrocytes (Figure 8C) (89). As shown in Figure 8D, the RNA velocity vector field for our four samples overlaid on the UMAP indicates that MH and FL mature oligo velocity vectors are shorter and pointing away from OPCs and pODs indicating the cells are in homeostasis. In contrast, the vectors in the FH and ML mature oligodendrocytes are longer indicating they are still undergoing rapid differentiation and many of the vectors are pointing away from the mature oligodendrocytes cluster and in many instances towards the pOD cluster which suggests many of the cells are still pre-myelinating oligodendrocytes (Figure 8D). Furthermore, all the vectors in the FH pODs are long and are pointing towards the OPCs and away from the mature oligodendrocytes indicating most cells in that cluster are still OPC-like (Figure 8D). Interestingly, the velocity vectors in the OPC cluster of all four samples are short in length suggesting OPCs are quiescent and that the lack of mature oligodendrocytes in the ML and FH is not due to a lack of OPCs (Figure 8D). Overall, RNA velocity vector field analysis suggests a sequential commitment from OPCs to pODs and then finally mature oligodendrocytes in FL and MH, while there appears to be very few mature oligodendrocytes in ML and FH which may be due to disruption of pOD differentiation. In Figure 8E, latent time analysis of the three clusters further suggests that there are very few terminally differentiated mature oligodendrocytes in ML and FH compared to FL and MH, while many OPCs in each sample are quiescent.

**Figure 8.**
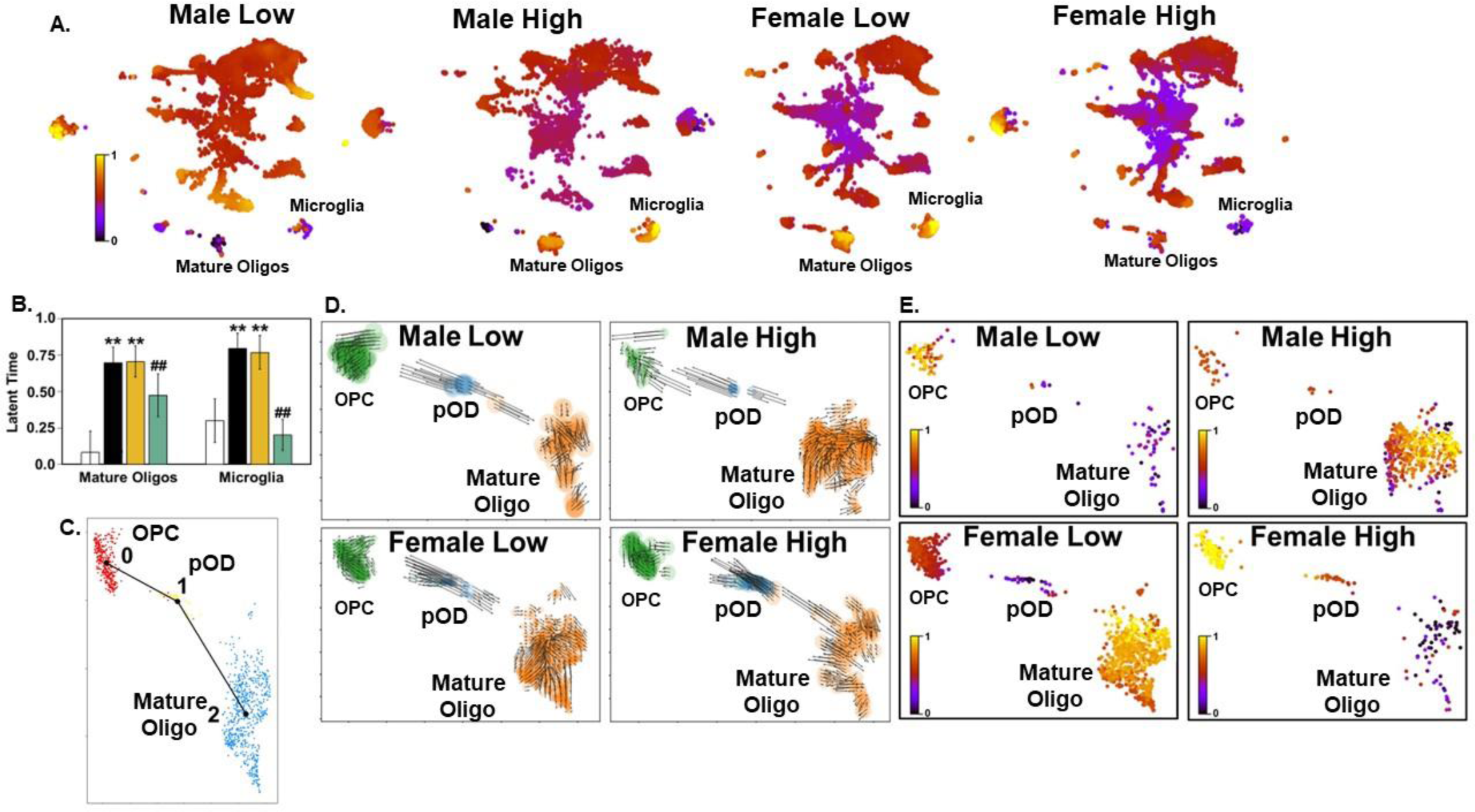
RNA velocity analysis reveals different developmental trajectories in OPCs and oligodendrocytes from C58/J amygdala that are dependent on mouse sex and sociability. (A) RNA velocity analysis latent time for Male Low Social (ML), Male High Social (MH), Female Low Social (FL), and Female High Social (FH) amygdala cell types projected on to a UMAP plot. (B) Graph of latent time in mature oligodendrocytes and microglia from ML, MH, FL, and FH amygdala. (C) Diagram of predicted developmental dynamics of OPCs, pre-oligodendrocytes (pOD), and mature oligodendrocytes where a cell differentiation trend moves from OPCs (0 or initial cluster) to pre-oligodendrocytes (1) to mature oligodendrocytes (2). (D) RNA velocities of OPCs, pre-oligodendrocytes, and mature oligodendrocytes from ML, MH, FL, and FH amygdala with arrows showing direction of development. (E) RNA velocity analysis latent time for OPCs, pre-oligodendrocytes, and mature oligodendrocytes from ML, MH, FL, and FH amygdala. Results reported as mean ± SEM (n = 1 mouse per group), **p<0.01 vs. ML, ##p<0.01 vs. FL & MH using Student’s t-test.

RNA velocity analysis also enabled us to investigate the transcriptional dynamics of driver genes during cellular state transition. Supplemental Figure 9 shows heatmaps of specific lineage driver genes whose relative expression levels are ordered along latent time. In Supplemental Figure 9C, the gene trend plots in FL follow the predicted expression trajectory of cluster-specific marker genes for OPCs (*Gria4*, *Plpp4*) that eventually differentiate into oligodendrocytes (*Rffl*). In contrast, the predicted expression trajectory of cluster-specific markers is reversed in ML and FH (Supplemental Figure 9A,D). For instance, in FH the latent time shows OPCs have a longer latent time and are quiescent compared to mature oligodendrocytes which have a shorter latent time (Supplemental Figure 9D). As a result of this, the early driver genes like *Bcas1os2* (marker of early myelinating oligodendrocytes), *Enpp6* (early marker of oligodendrocyte differentiation), and *Pld1* (marker of the morphological differentiation of oligodendrocytes prior to myelination) are associated with mature oligodendrocytes and differentiating oligodendrocytes rather than OPCs (Supplemental Figure 9D).

## Discussion

Our comparison of the C57BL/6J and C58/J inbred strains identified transcriptional and pathway signatures that suggest immune-related biological processes differ in amygdala cells from C58/J, a model of ASD-like behavior. We identified differentially hyper-and hypomethylated regions in genes in C58/J amygdala when compared to C57BL/6J amygdala. snRNA-Seq data from the C58/J amygdala identified a consistent transcriptional signature in mature oligodendrocytes and microglia characterized by altered ASD risk gene expression and impaired expression of myelin-related genes and microglial homeostatic genes that was dependent on sex and sociability. Alterations in gene regulatory networks, impaired cell-to-cell communication, and aberrant cell differentiation were identified in microglia and mature oligodendrocytes that may help explain why a reduced density of both cell types was observed in ML and FH C58/J amygdala. Many of these features were verified in our bulk RNA-Seq data from C58/J amygdala. Taken together, these results provide preclinical evidence that defects in mature oligodendrocytes and microglia play an important role in sociability which is one of the core deficits of ASD.

### The C58/J mouse model is a useful preclinical tool for understanding the mechanistic basis of sociability deficits

The C58/J strain has several behavioral features that reflect symptoms of ASD, including a lack of social preference in the 3-chamber social choice test, hyperactivity, and abnormal repetitive responses (37, 90-92). A previous study has shown that in contrast to C57BL/6J mice, C58/J and other low sociability strains display divergent phenotypes, such that approximately 50% of mice have positive sociability while 50% show marked social avoidance (36). This provides an opportunity to investigate both genetic risk for ASD-like social deficits, through a between-strain comparison with C57BL/6J, and risk based on epigenetic changes, by within-strain comparisons between high sociability and low sociability isogenetic C58J mice.

In our studies, bulk RNA-Seq analysis of amygdala from C58/J and C57BL/6J mice revealed immune-related genes and pathways were downregulated in C58/J amygdala compared to C57BL/6J. Alterations in the immune system are a major factor contributing to the pathogenesis of ASD (65). Our data suggests differentially altered immune system pathways including IL-1β, TNFα, adaptive immunity, MAPK signaling, and T-cell immunity which have been shown to be dysregulated in ASD (65, 93). Several immune-related genes were differentially expressed between the strains that play a role in ASD including *Pla2* genes, innate immunity genes, and the axon guidance molecule *Robo4*. In addition to bulk RNA-Seq analysis, our differential methylation analysis suggests C58/J and C57BL/6J mice have distinct epigenetic profiles that indicate distinct epigenetic signatures between high social and ASD-like low social mice that may contribute to sociability differences. Overall, the between-strain comparison indicates that C58/J has transcriptional profiles with significant alterations in multiple genes implicated in the pathogenesis of ASD.

Our snRNA-Seq analysis of the C58/J amygdala provided confirmation of our bulk RNA-Seq data and identified the ML group as having the most ASD risk gene changes in line with higher prevalence of ASD diagnoses in males. The ML differed from both the MH (based on sociability) and the FL (based on sex) via upregulation of SFARI ASD risk genes as well as ASD-related pathways in multiple glutamatergic, GABAergic, and non-neuronal cell types. Cell type specific analysis showed that mature oligodendrocytes and microglia in ML exhibited the greatest number of differentially expressed SFARI ASD risk genes compared to the other cell types. Similar to across-strain changes identified by our comparison of C58/J and C57BL/6J, ML microglia exhibited down regulation of several immune-related pathways including Toll-Like receptor 2 signaling, cytokine production, MAPK signaling, myeloid cell activation, and microglial activation in comparison to the MH and FL groups. These results are significant because microglia are involved in synaptic pruning and neuronal development, while also promoting the formation of new synapses, supporting myelination/oligodendrogenesis, and supporting cell health (94). Our data indicate microglia cell number was reduced in ML compared to MH and FL. RNA velocity analysis indicated aberrant microglia differentiation in ML, while cell-to-cell interaction analysis indicated upregulated Ephrin signaling which is known to stimulate cell differentiation (85). The reduced numbers of ML microglia could contribute to altered immune signaling and would have negative implications for synaptic pruning, synapse formation, and cell clearance. Further, the reduced number of microglia and altered pathway expression may have additional effects on oligodendrogenesis and myelination (94).

### Myelination defects in the C58/J mouse model

Multiple mouse models of ASD have reported impaired myelination. Tuberous sclerosis complex 1 (*Tsc1*) knockout mice display hypomyelination and reduced oligodendrocyte density (95), while fragile X (Fmr1) knockout mice display delayed myelination and reduced OPC density in cerebellum (96). Mouse mutants for *Cntnap2* also exhibit delayed myelination (97). Our snRNA-Seq studies revealed mature oligodendrocytes from ML amygdala exhibit downregulation of myelin-related genes and pathways compared to MH (sociability) and FL (sex). ML mature oligodendrocytes also exhibited reduced cell number compared to FL and MH mature oligodendrocytes. RNA velocity analysis and cell-to-cell interaction analysis revealed differentiation is impaired in ML mature oligodendrocytes and several pathways that influence oligodendrocyte differentiation were altered in ML including Bmp, vitronectin, and contactin signaling. Many of these pathways in ML involved pericyte signaling to OPCs and mature oligodendrocytes and studies suggest pericytes influence OPCs via physical and functional interactions (98). GRN analysis revealed an absence of *Sox8* and *Sox10* regulon activity in ML which appear to drive expression of DEGs that promote myelin formation in MH and FL mature oligodendrocytes.

*Tcf4* mutant mice are an ASD mouse model with oligodendrocytes that display a similar profile compared to C58/J ML and FH mature oligodendrocytes (99). Specifically, mutation in the *Tcf4* gene resulted in reductions in genes involved in myelin formation, reductions in mature oligodendrocyte numbers, but no change in OPC density (99). Single cell analysis of human control and ASD cells from cortical brain tissue have previously revealed reduced mature oligodendrocytes and microglial cell density in ASD samples while no change in OPC cell density was found (99, 100). Interestingly, TEM imaging showed *Tcf4* mutant mice display reduced mature oligodendrocytes in the corpus callosum as well as a reduction in myelinated axons (99). Several other neuroimaging studies have also reported white matter deficiency in ASD patients (101). However, our MRH analysis revealed no differences in connectomes or brain volumes between our male and female low and high social C58/J brains. It is important to note that hypomyelination and reduced mature oligodendrocyte density is not a universal finding in mouse ASD models. *Pten* knockout mice display hypermyelination in brain, while knockdown of *MeCP2* in primary rat oligodendrocytes increased synthesis of genes associated with myelin synthesis (*Plp1*, *Mog*, *Mbp*) (102, 103). Studies have also shown that ASD-related myelination deficits are region-specific and that some brains from the same human ASD patient have shown opposite changes in myelination in different brain regions such as prefrontal cortex and cerebellum (104). What is clear, however, is that altered oligodendrocyte biology and myelination play an important role in ASD.

### An ASD-like, myelin-deficient, inflammatory transcriptomic profile in FH C58/J

A surprising finding in our studies was that an ASD-like transcriptome profile was discovered in FH mature oligodendrocytes and microglia. Given that ASD prevalence is elevated in males and is defined by poor social skills and functioning, one would predict MH (sex) and FL (sociability) mature oligodendrocytes and microglia would look more ASD-like when compared to those same cell types in FH. Our data revealed that this was not the case for our analysis of C58/J. In support, our bulk RNA-Seq analysis revealed significant differences in ASD-related biological pathways between MH and FH amygdala. Moreover, the bulk RNA-Seq data confirmed that MH amygdala exhibited upregulation of pathways related to myelin assembly and axon ensheathment when compared to FH amygdala. This finding validated our discovery that FH mature oligodendrocytes exhibit downregulated pathways associated with myelination and axon ensheathment compared to MH mature oligodendrocytes.

Interestingly, our bulk RNA-Seq data also showed oxytocin administration upregulated biological pathways related to myelin assembly, axon ensheathment, and oligodendrocyte differentiation in FH amygdala. This action could have been the basis for the prosocial oxytocin effects previously reported in C58/J and other mouse models (38, 39). These data also provide potential insight into the mechanisms by which oxytocin, as a potential ASD therapeutic, might function with respect to rectifying deficiencies in myelin-related pathways and signaling in mature oligodendrocytes within the FH amygdala. Leading edge analysis of bulk RNA-Seq data in FH oxytocin-treated amygdala also identified several genes (e.g., *Opalin*) whose expression was increased in FH oxytocin-treated amygdala that play a role in oligodendrocyte differentiation (Supplemental Figure 10B). This finding supports our cell-to-cell interaction data and RNA velocity data that suggest oligodendrocyte differentiation is impaired in FH mature oligodendrocytes which contributes to reduced FH mature oligodendrocyte cell density. In addition to aberrant myelination, the FH is also interesting because of its inflammatory profile. Bulk RNA-Seq data showed genes (*Tlr1*, *Il15ra*, *Edn1*) and pathways were upregulated in FH amygdala compared to MH amygdala that are associated with an inflammatory response (Supplemental Figure 11B-D), while oxytocin treatment also decreased pathways associated with detoxification of reactive oxygen species and oxidative stress in FH amygdala (Supplemental Figure 10A). These findings in the bulk RNA-Seq data support snRNA-Seq data in FH that pericytes signal to microglia via the proinflammatory Spp1 pathway. Our findings in FH raise questions about sex-specific differences in clinical presentation and diagnosis between males and females with ASD. Our findings that ML and FH samples are the most ASD-like suggests ASD symptoms and presentations may be different in males and females regarding at least one aspect of typical ASD core deficits like sociability and this may be why females are underdiagnosed (105). One possibility is that amygdala plays an inhibitory role to limit risky or impulsive social approach to unfamiliar strangers, and alterations in amygdala function could lead to abnormal hypersociability in normally cautious female mice.

### Limitations

Our findings are significant. However, several caveats can be noted. We have only investigated a single model of ASD-like behavior, and a single aspect (social approach) of the complex behavior domain. Further work is needed to determine whether our findings generalize to a broader range of mouse models of genetic and environmental ASD risk, whether similar epigenetic signatures in brain underlie other social phenotypes, and the contribution of brain regions, other than amygdala, to ASD-like social deficits. The cellular mechanisms linking reduced oligodendrocyte differentiation and reduced myelination to an ASD phenotype in C58/J mice needs further investigation. Additional snRNA-Seq studies would be needed to determine if effects in oligodendrocytes and microglia are unique to amygdala or if this occurs in other brain regions. In our study, the subchronic ocytocin regimen did not shift the pattern of social divergence in the C58/J model (Supplemental Figure 1A-D). Thus, the effects of oxytocin need further examination to better understand its potential as an ASD therapeutic. Cell proportion estimates may differ between ML, MH, FL, and FH amygdala in snRNA-Seq experiments due to slight differences in where the tissue originated when dissected.

## Conclusions

In summary, our work demonstrates the utility of the C58/J mouse model in evaluating the influence of sex and sociability on the transcriptome in brain regions that play an important role in ASD. Our single-nucleus transcriptome analysis elucidates the pathological roles of oligodendrocytes and microglia from amygdala in ASD. Our study provides many details regarding specific regulatory features that are disrupted in these two cell types that include transcriptional gene dysregulation (ASD risk genes, myelinating genes, microglial homeostatic genes), aberrant cell differentiation, impaired gene regulatory networks, and alterations in key pathways that promote microglia and oligodendrocyte differentiation. Our work also reveals the cellular changes oxytocin administration produces within cells of the brain. Future research is needed to determine if targeting oligodendrocytes and microglia could be developed into effective treatments for ASD. Research at the single cell level should certainly provide more potential ASD therapeutic targets.

## List of Abbreviations

ASD: Autism Spectrum Disorder
Bulk RNA-Seq: Bulk RNA-Sequencing
DEG: Differentially Expressed Gene
FH: Female High Social
FL: Female Low Social
GRN: Gene Regulatory Network
MH: Male High Social
ML: Male Low Social
snRNA-Seq: single nucleus RNA-Sequencing

## Declarations

### Ethics approval

All experimental procedures were conducted in compliance with an approved UNC IACUC protocol, and those set forth in the “Guide for the Care and Use of Laboratory Animals” as published by the National Research Council.

### Consent for publication

Not applicable

### Availability of data and materials

The datasets used and analyzed during the current study are available from the corresponding author on reasonable request.

### Author’s Contributions

Conceptualization, S.G.G., S.S.M., and Y.H.J.; Methodology, Y.H.J., S.S.M., and S.G.G.; Formal Analysis, Investigation, Visualization, G.D.D., S.K.S., V.D.N., G.P.C., K.H., Y.Q., G.A.J., S.S.M. and S.G.G.; Writing – Original Draft, G.D.D., G.A.J., S.G.G., S.S.M., and Y.H.J.; Writing – Review & Editing, G.D.D., S.G.G., S.S.M., Y.H.J.; Supervision, S.G.G., S.S.M. and Y.H.J.; Funding Acquisition, S.G.G., S.S.M. and Y.H.J.

## Acknowledgements

We would like to thank Dr. Devjanee (Devi) Swain Lenz and the rest of the staff at the Duke Sequencing and Genomic Technologies (SGT) Core, as well as the staff at the Duke Molecular Genomics Core, particularly Karen Abramson, Stephanie Arvai, and Emily Hocke for their excellent support throughout the course of this study. We also thank the staff at the Mouse Behavioral Phenotyping Laboratory, Carolina Institute for Developmental Disabilities, at the University of North Carolina at Chapel Hill.

## Figure Legends

**Supplemental Figure 1.**
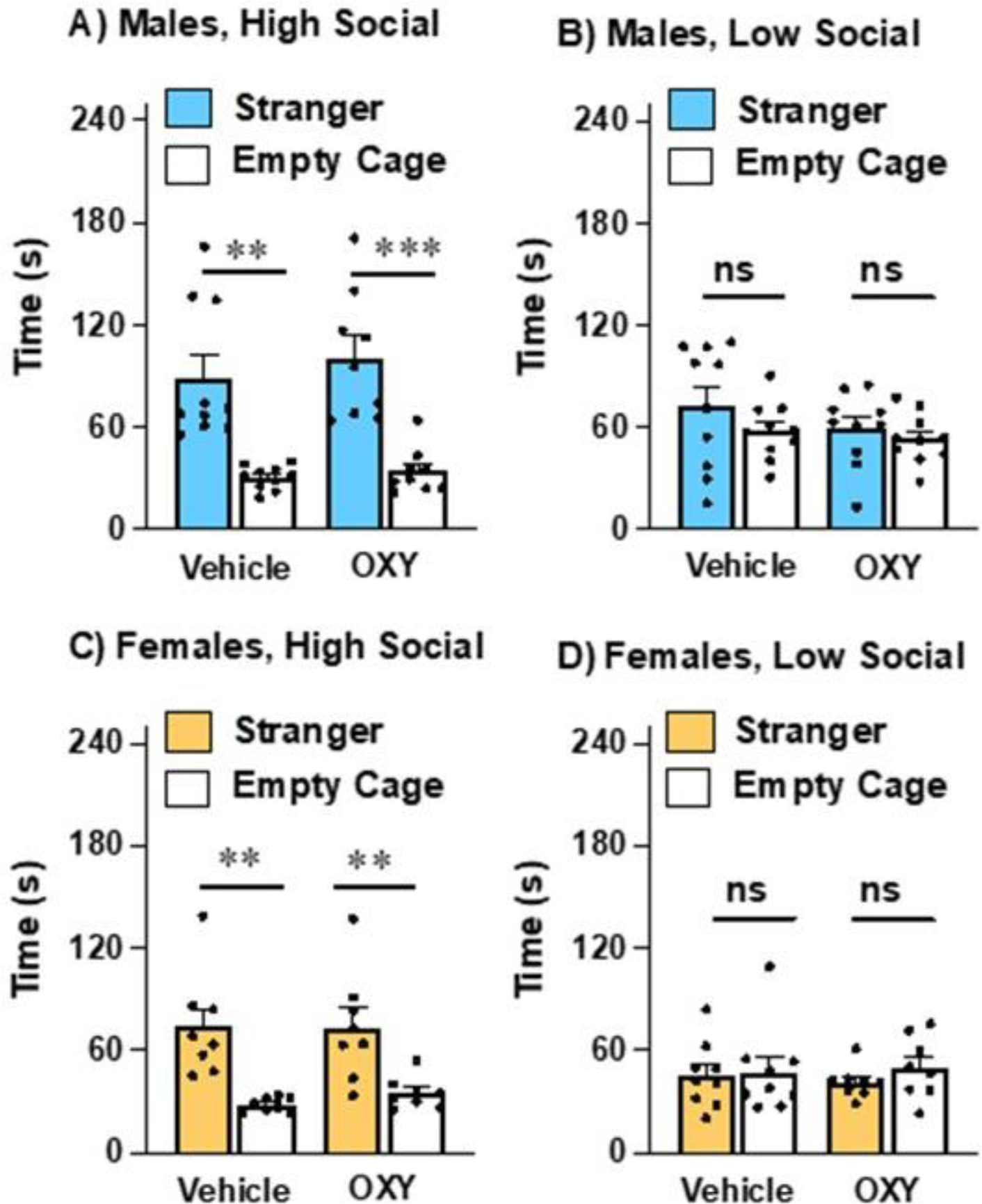
Lack of oxytocin effects on divergent phenotypes in C58/J mice. (A-D) Comparison of social preference based on time spent in close proximity to a stranger mouse during the 3-chamber social choice test in male and female C58/J high social and low social mice that received vehicle or oxytocin (OXY). Data are mean ± SEM time spent within 5.0 cm proximity to a cage containing a stranger mouse, or an empty cage. N=8-10 mice per group. *p<0.05, **p<0.01, ***p<0.001, ns (nonsignificant); within-group repeated measures comparison, of side.

**Supplemental Figure 2.**
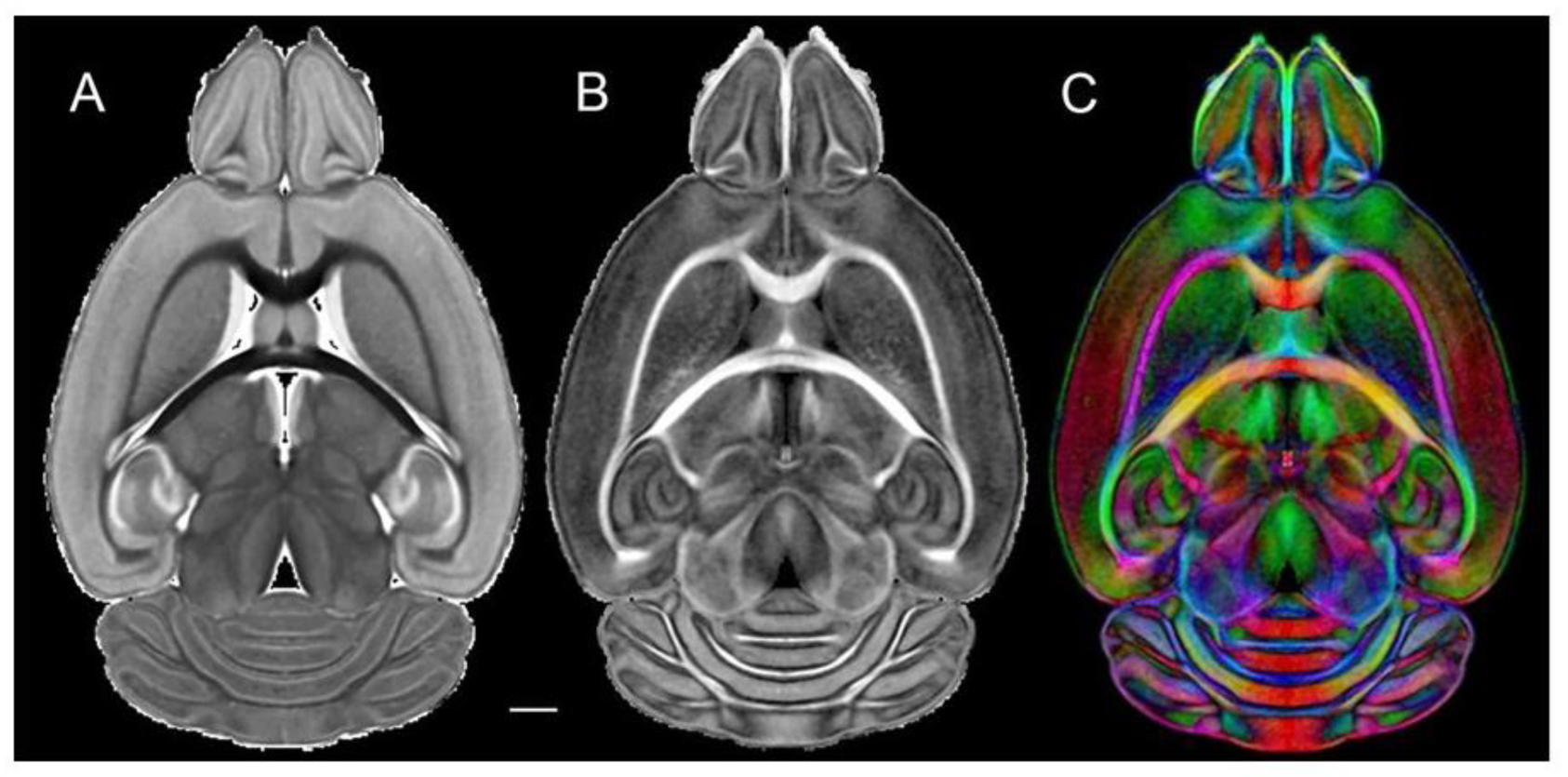
Images produced from Magnetic Resonance Histology (MRH) of C58/J mouse brain. The MRH processing pipeline produced a series of images in which the signal highlights different diffusion parameters and brain anatomy. The figure shows representative images from the average of a C58/J Male High Social mouse where Image A shows Mean Diffusivity (MD), Image B shows fractional anisotropy (FA), and Image C shows color fractional anisotropy (cIrFA). The scale bar is 1 mm.

**Supplemental Figure 3.**
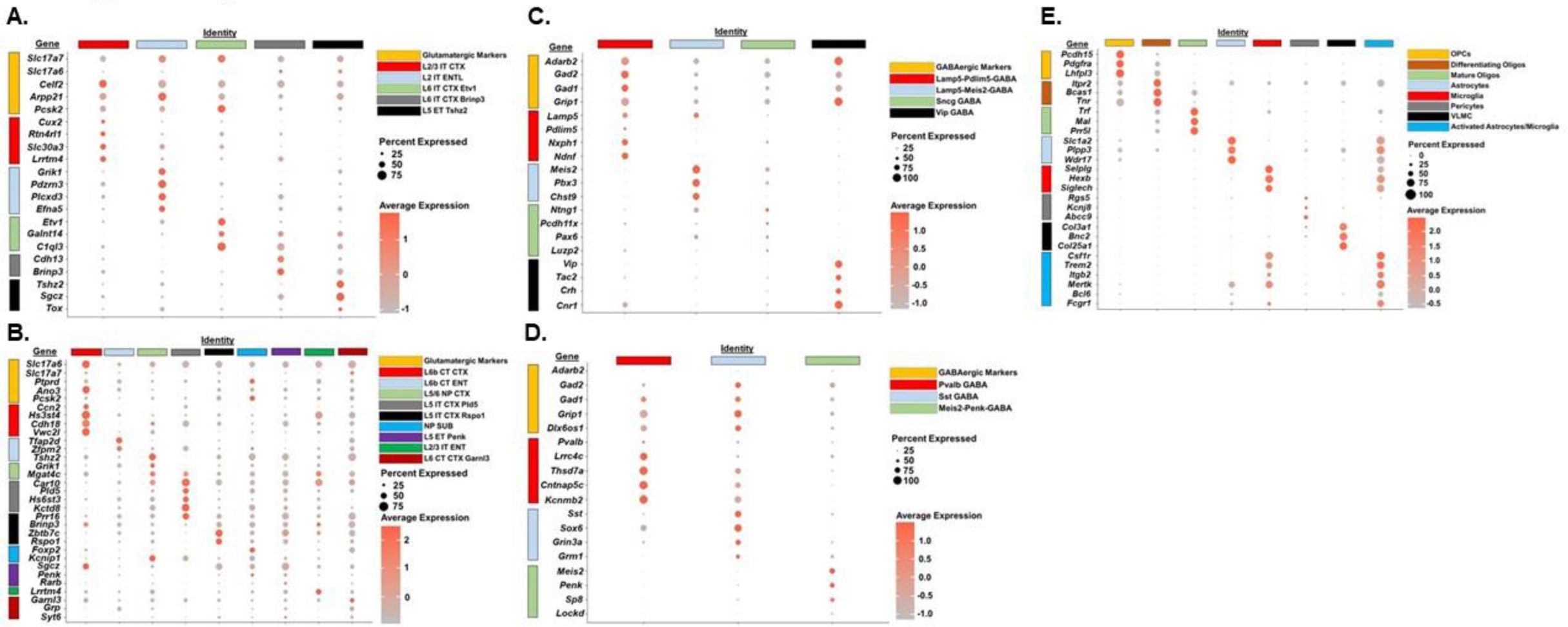
Glutamatergic, GABAergic, and Non-Neuronal cell markers in C58/J mouse amygdala. Dotplots of (A,B) Glutamatergic, (C,D) GABAergic, and (E) Non-Neuronal cell clusters showing percentage of cells within each cluster derived from amygdala taken from C58/J male and female mice and marker gene expression in each cluster. Dot size indicates percentage of cells within a cluster, while color shows average expression level across all cells within a cluster.

**Supplemental Figure 4.**
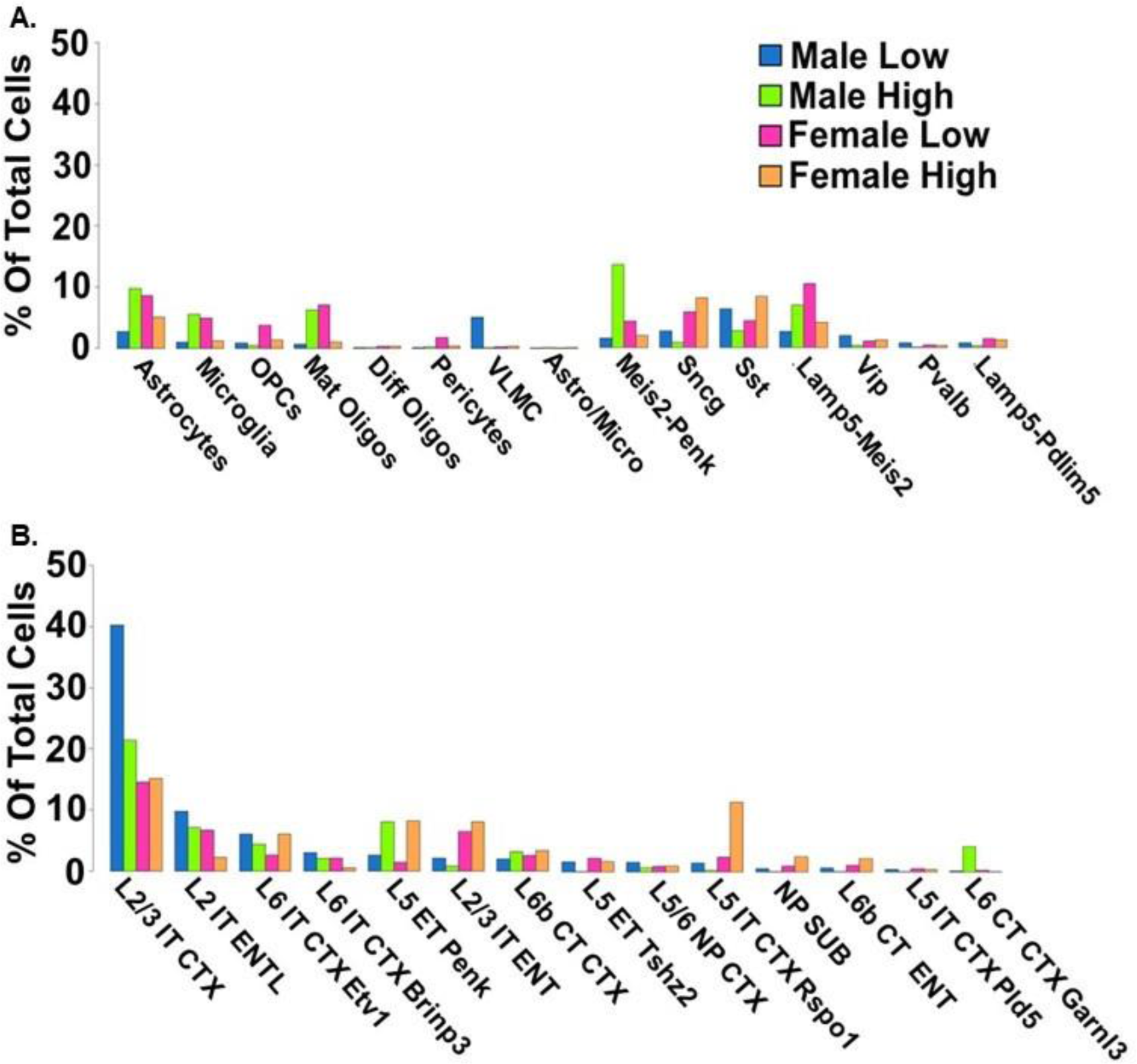
Percentage of 29 cell types across C58/J amygdala depends on sex and sociability. Percentage of (A) non-neuronal and GABAergic and (B) glutamatergic cell types in Male Low Social, Male High Social, Female Low Social, and Female High Social amygdala (n = 1 mouse per group).

**Supplemental Figure 5.**
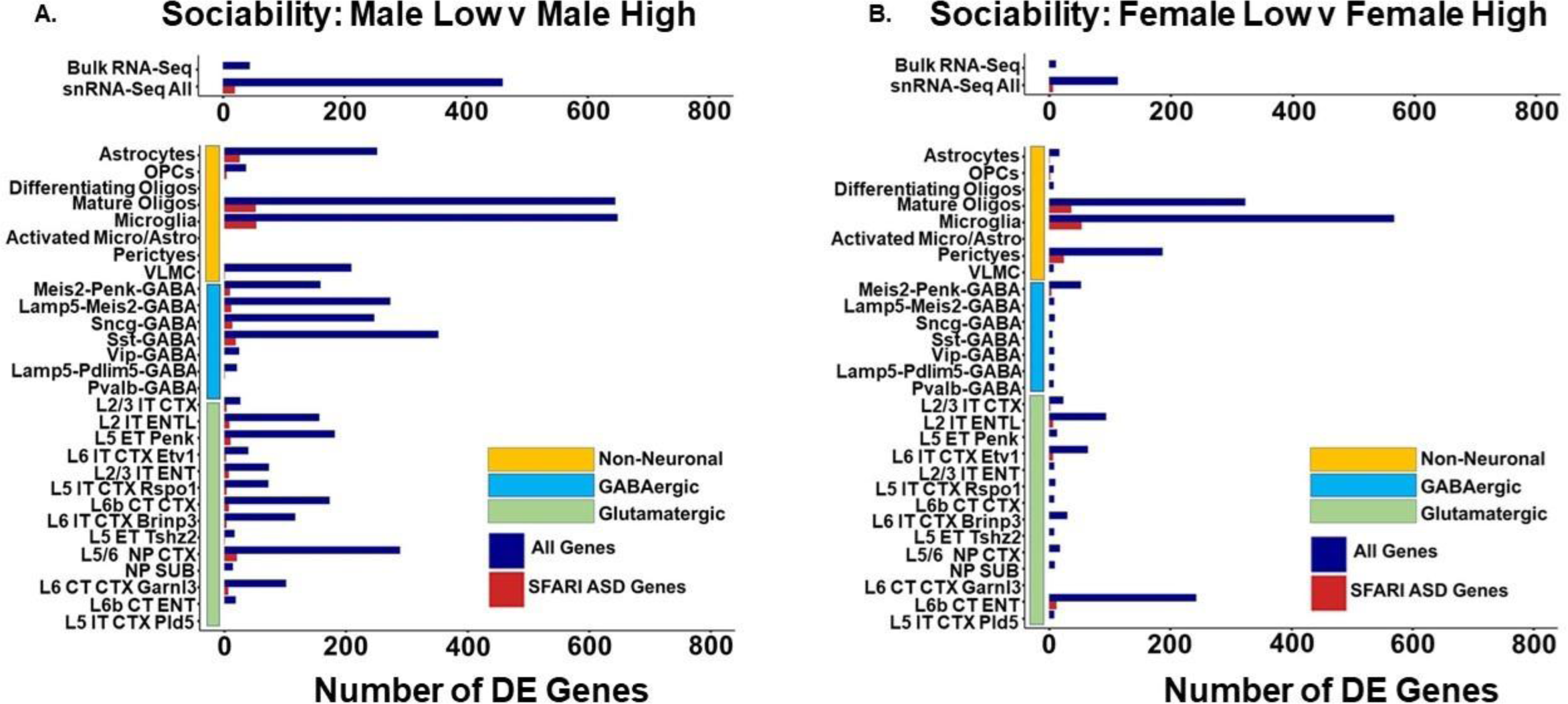
Effect of sociability on differential gene expression in C58/J amygdala. (A-B) Bar graphs showing total differentially expressed genes (DEGs) and SFARI ASD risk DEGs in whole amygdala and in individual cell types when C58/J mouse groups were compared based on sociability (n = 1 mouse per group). DEGs in whole amygdala were obtained via bulk RNA-Seq, while DEGs in individual cell types were obtained via snRNA-Seq.

**Supplemental Figure 6.**
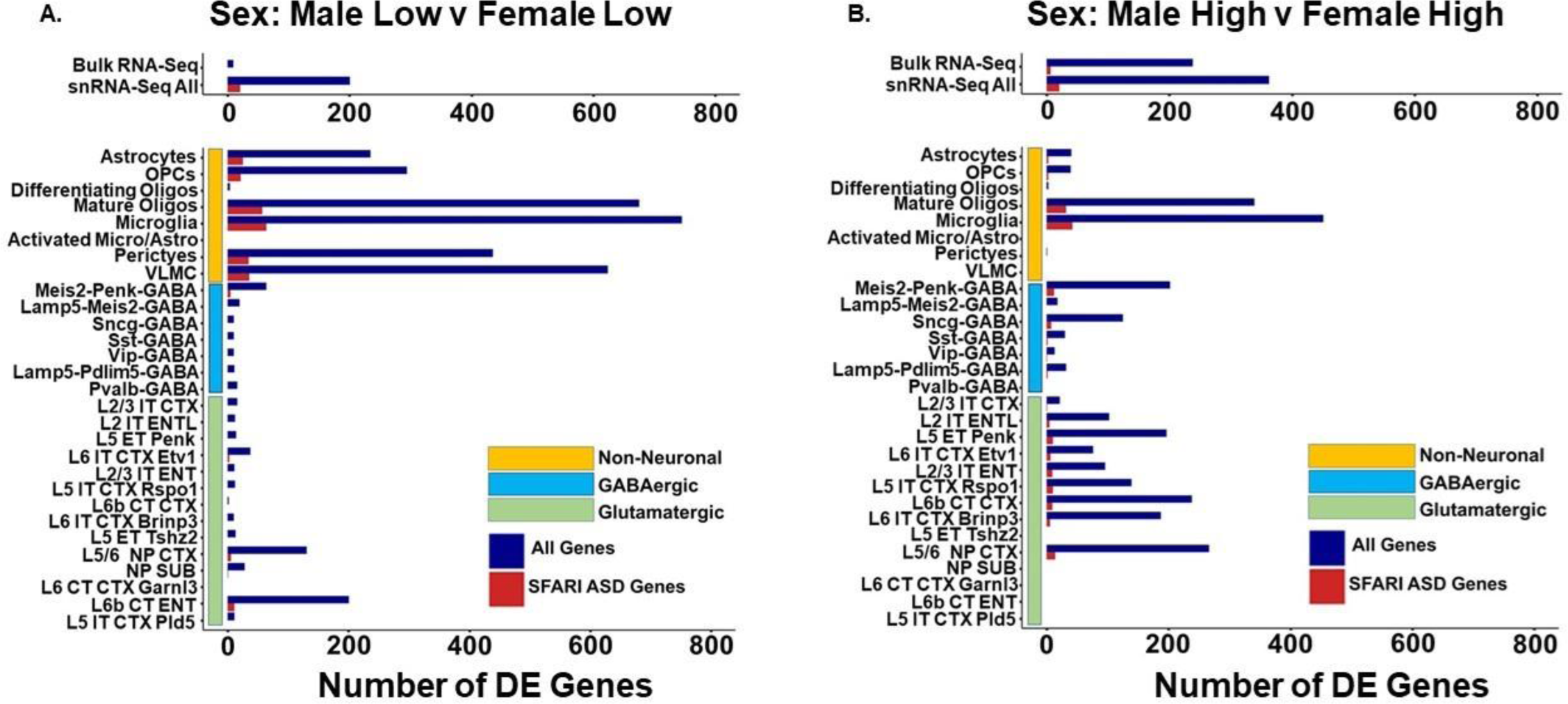
Effect of sex on differential gene expression in C58/J amygdala. (A-B) Bar graphs showing total differentially expressed genes (DEGs) and SFARI ASD risk DEGs in whole amygdala and in individual cell types when C58/J mouse groups were compared based on sex (n = 1 mouse per group). DEGs in whole amygdala were obtained via bulk RNA-Seq, while DEGs in individual cell types were obtained via snRNA- Seq.

**Supplemental Figure 7.**
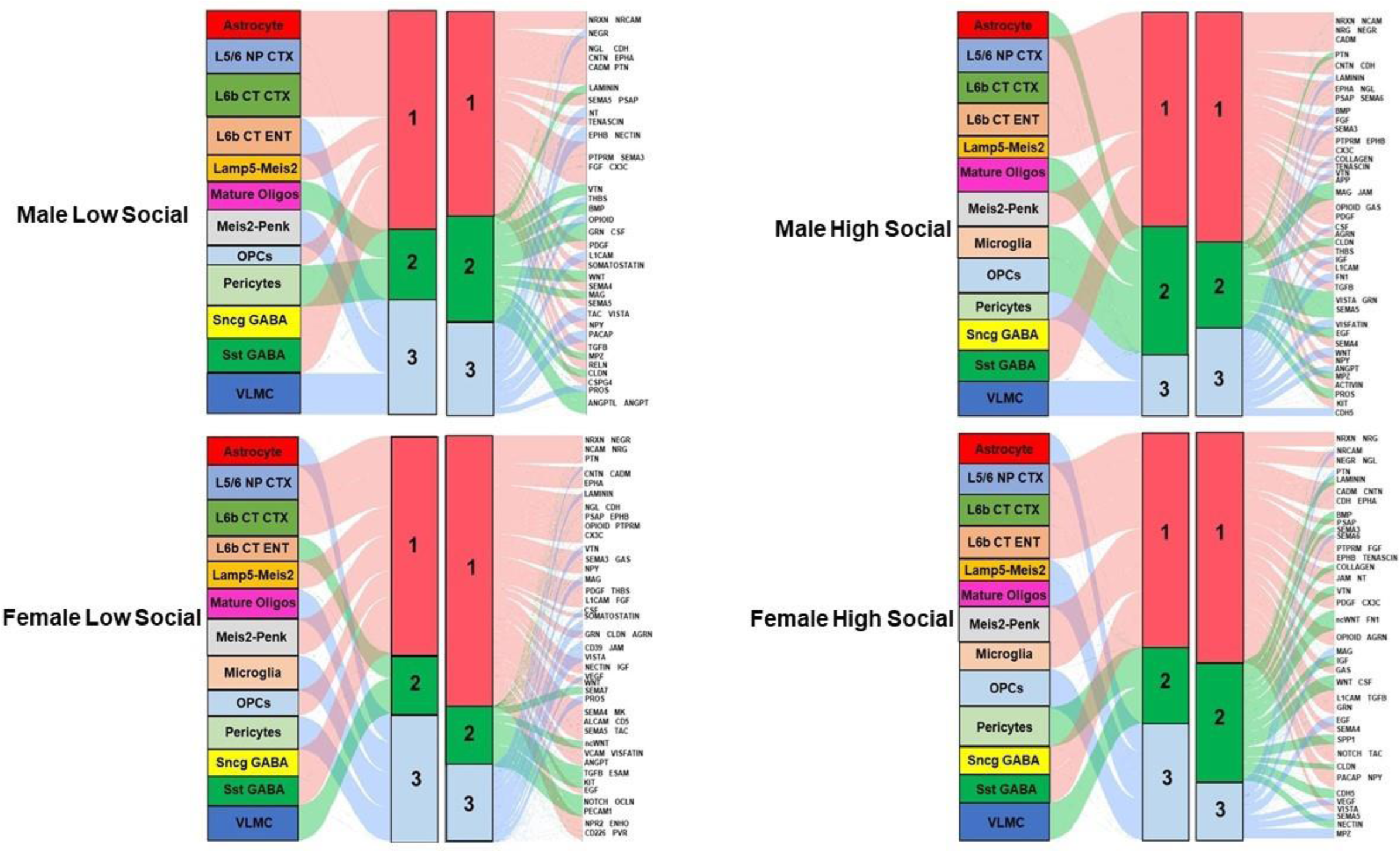
Patterns of outgoing signaling from select cell types to OPCs and mature oligodendrocytes in C58/J mouse amygdala. (A-D) The outgoing signaling patterns of secreting cells in amygdala to OPCs and mature oligodendrocytes are visualized by alluvial plot and show the correspondence between the inferred latent signaling patterns (labeled 1, 2, and 3) and specific cell groups, as well as specific signaling pathways. The thickness of the flow indicates the contribution of each cell group or signaling pathway to each latent pattern. The height of each pattern is proportional to the number of its associated cell groups or signaling pathways.

**Supplemental Figure 8.**
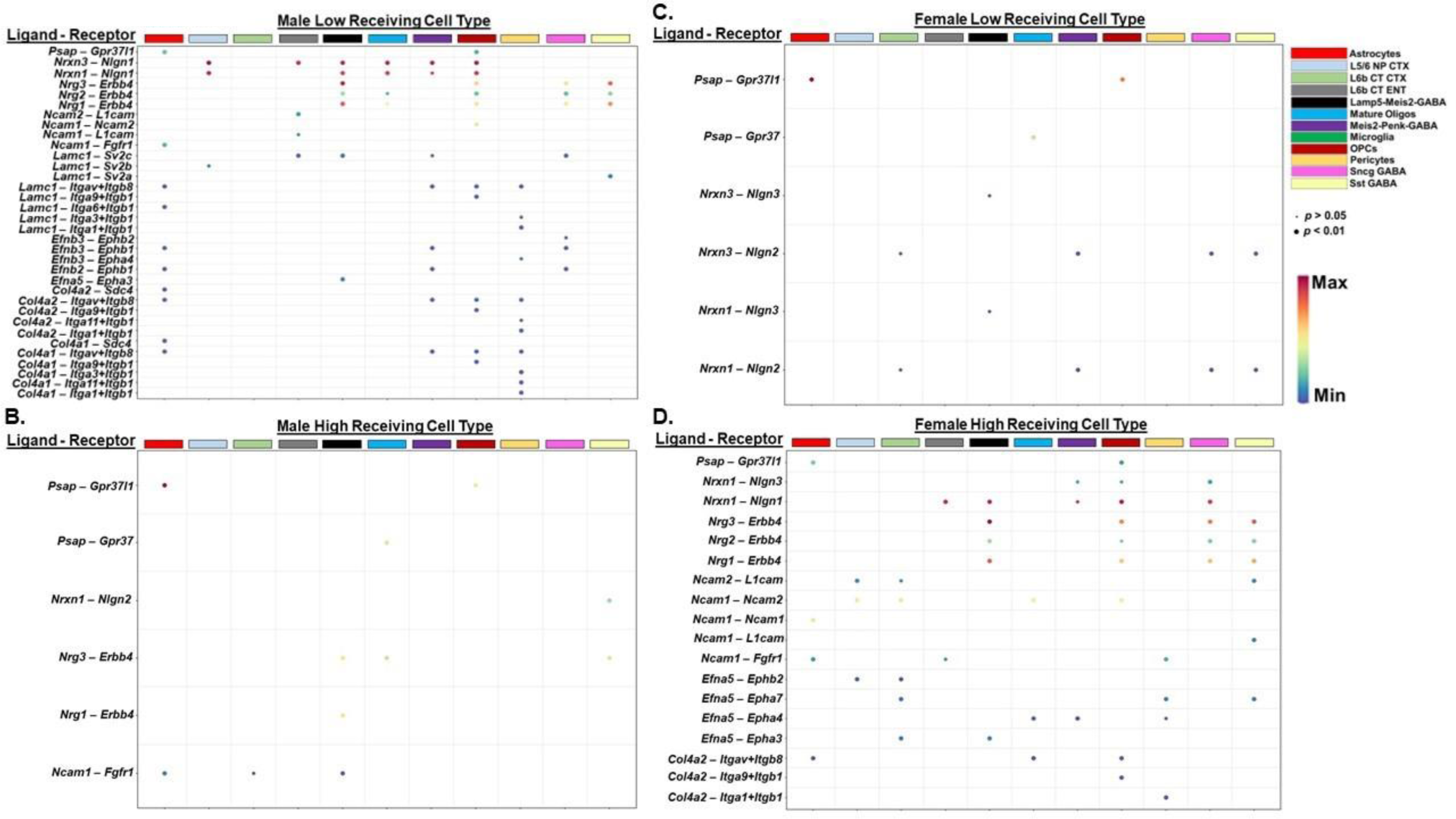
Patterns of outgoing signaling from microglia to select cell types in C58/J mouse amygdala. (A-D) Dot plots showing specific unidirectional signaling originating from microglia which releases specific ligands that bind to receptors on the indicated cell types in representative groups from C58/J mouse amygdala (n = 1 mouse per group). Color implies expression magnitude and dot size indicates specificity of the interaction.

**Supplemental Figure 9.**
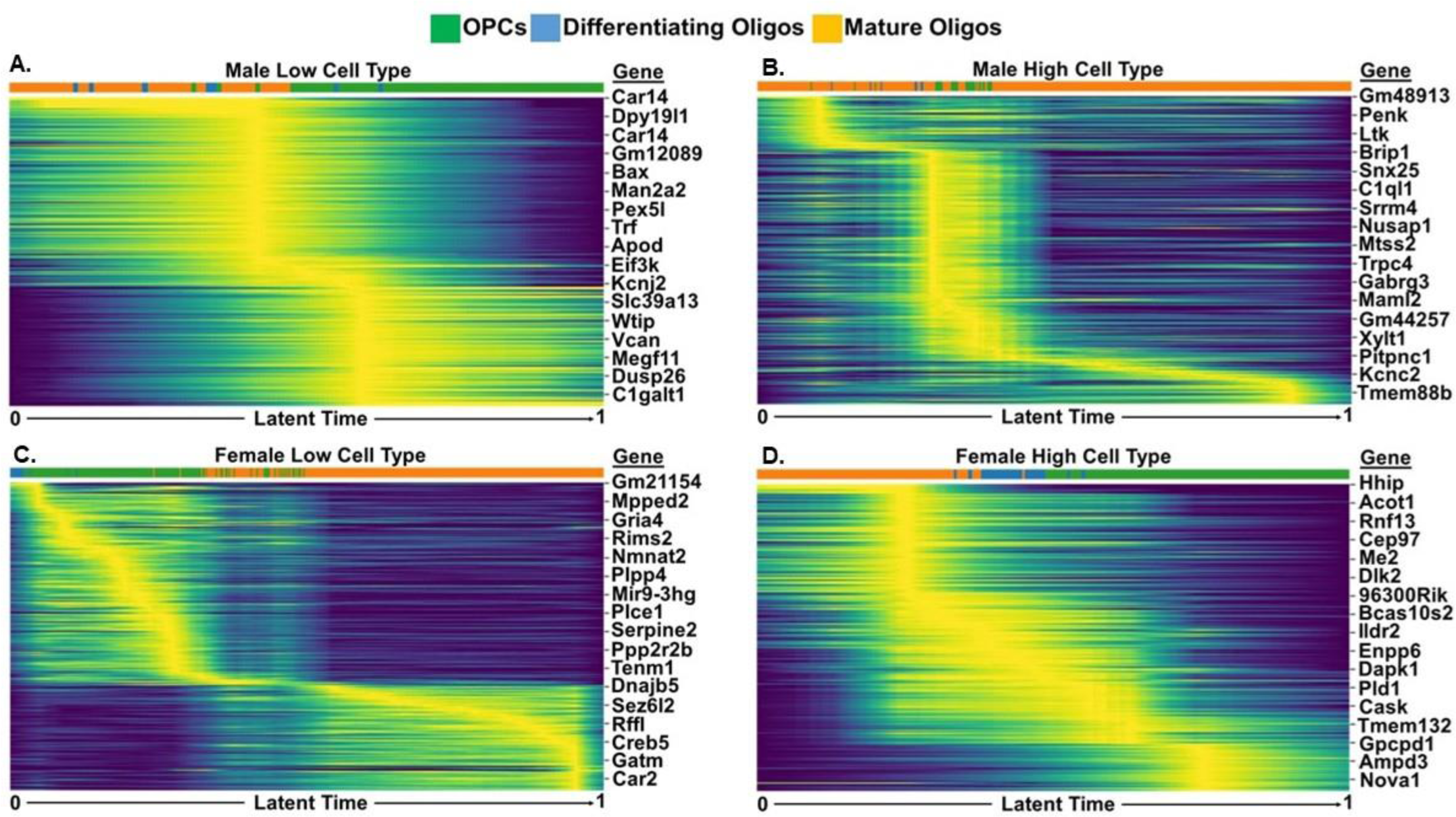
Heatmap of gradual gene expression changes along the latent time trajectory in C58/J amygdala OPCs, differentiating oligodendrocytes, and mature oligodendrocytes. Each column shows gene expression levels in the indicated cell type at each point in latent time and each row represents a corresponding gene in (A) Male Low Social, (B) Male High Social, (C) Female Low Social, and (D) Female High Social amygdala (n = 1 mouse per group). The gradient from blue to yellow indicates the amount of gene expression from lowest to highest.

**Supplemental Figure 10.**
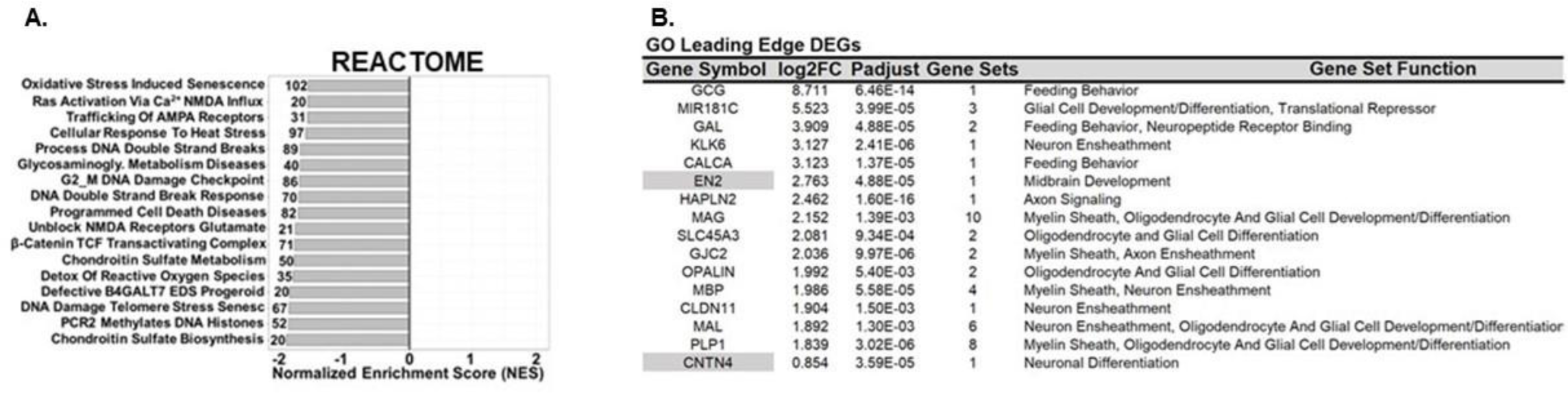
GSEA identifies differentially expressed REACTOME pathways from bulk RNA-Seq data generated in Female High Social Vehicle amygdala when compared to Female High Social Oxytocin amygdala. (A) Reactome enrichment analysis of pathways in Female High Social Vehicle (FHV) amygdala compared to Female High Social (FHOXY) amygdala obtained by bulk RNA-Seq analysis and GSEA (n = 4-5 mice per group). Negative normalized enrichment score (NES) indicates pathways are upregulated in the FHV group. Numbers next to bars indicate total number of genes in pathway. (B) Differentially expressed genes that were identified as leading edge genes following GSEA analysis of bulk RNA-Seq data from FHV and FHOXY amygdala. Positive log2FC indicates genes were upregulated in FHOXY. Gene sets indicate number of gene sets that contain the DEG.

**Supplemental Figure 11.**
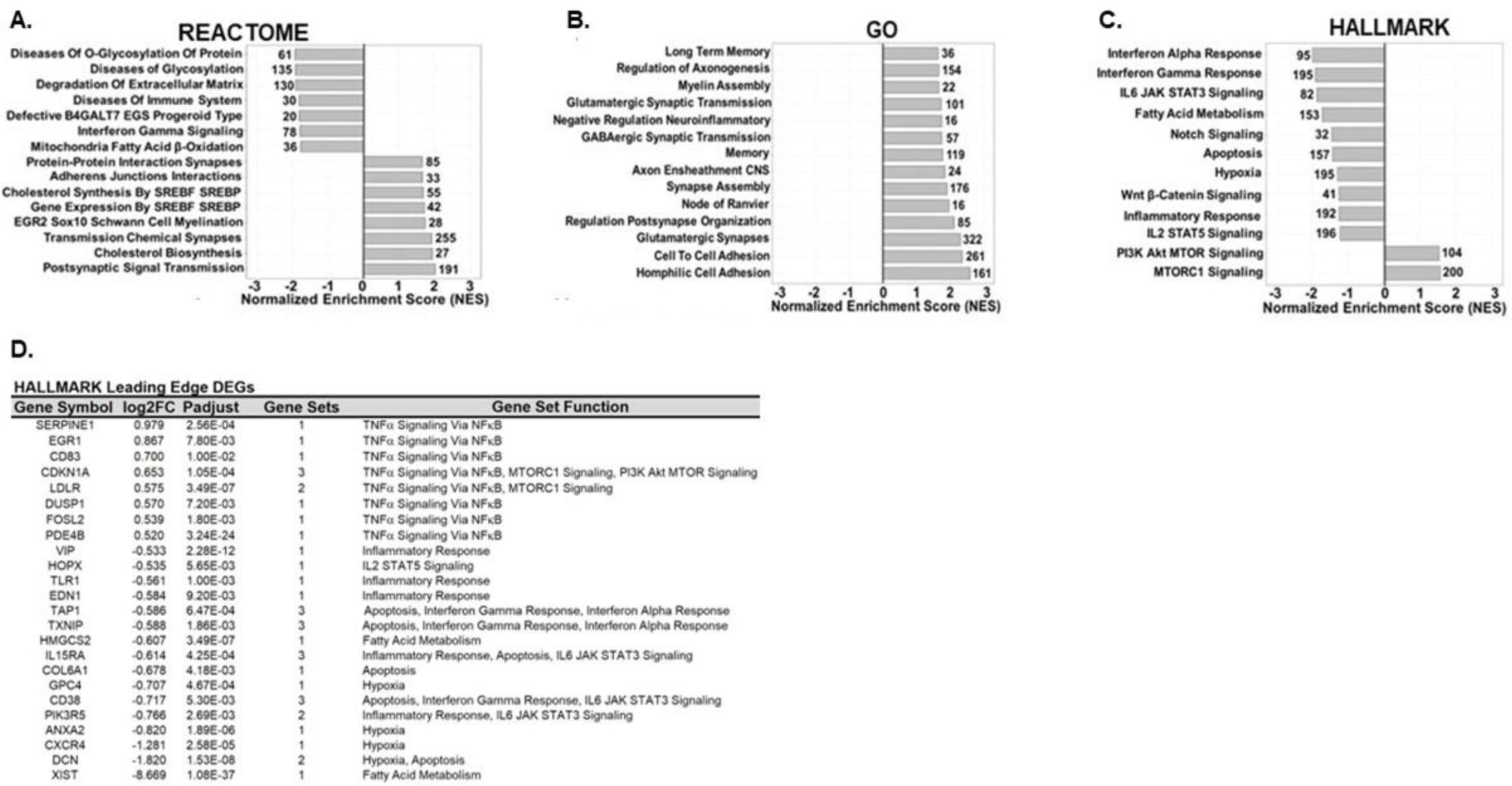
GSEA identifies differentially expressed REACTOME, GO, and HALLMARK pathways from bulk RNA-Seq data generated in Male High Social amygdala when compared to Female High Social amygdala. (A) Reactome, (B) Gene Ongology (GO), and (C) Hallmark enrichment analysis of pathways in Male High Social (MH) amygdala compared to Female High Social (FH) amygdala obtained by bulk RNA-Seq analysis and GSEA (n = 5 mice per group). Positive normalized enrichment score (NES) indicates pathways are upregulated in the MH group. Numbers next to bars indicate total number of genes in pathway. (D) Differentially expressed genes that were identified as Hallmark leading edge genes following GSEA analysis of bulk RNA-Seq data from MH and FH amygdala. Positive log2FC indicates genes were upregulated in MH. Gene sets indicate number of gene sets that contain the DEG.

